# Synthesising the multiple impacts of climatic variability on community responses to climate change

**DOI:** 10.1101/2021.04.26.441437

**Authors:** J. Christopher D. Terry, Jacob D. O’Sullivan, Axel G. Rossberg

## Abstract

Recent developments in understanding and predicting species responses to climate change have emphasised the importance of both environmental variability and consideration of the wider biotic community. However, to date, the interaction between the two has received less attention. Both theoretical and empirical results suggest that the combined effect of environmental variability and interspecific interactions can have strong impacts on existing range limits. Here we explore how competitive interactions and temporal variability can interact with the potential to strongly influence range shift dynamics. We highlight the need to understand these between-process interactions in order to predict how species will respond to global change. In particular, future research will need to move from evaluating possibilities to quantifying their impact. We emphasise the value and utility of empirically parameterised models to determine the direction and relative importance of these forces in natural systems.

## Introduction

Climate change is forcing species across the world to either adapt to different environments in-situ or shift their range to track moving climates. A signal of climate-change induced spatial displacement is clearly visible in shifts in the observed distribution of species across the globe (Parmesan and Yohe 2003, Lenoir et al. 2020). Improving our understanding of how ranges will shift is critical to future conservation efforts and ecosystem management (Pecl et al. 2017). Here we argue that multiple strands of ecological theory regarding the direct and indirect impacts of climate variability in determining community-level responses can be informative to this wider effort.

Underlying long-term climatic trends is higher-frequency environmental variation. This is partly cyclical (seasonal and diurnal) but there is also a considerable stochastic element. Differences in mean temperature between years are often comparable to decades of mean climate change (Huntingford et al. 2013). It is well established that environmental variability can have far-reaching impacts on populations (Coulson et al. 2004, Lawson et al. 2015, Boettiger 2018, Shoemaker et al. 2020a). On top of this, interactions with other species strongly influence a species’ range (Sexton et al. 2009, Kraft et al. 2015, Sirén and Morelli 2020). As much as climate driven range shifts are fundamentally driven by the dependence of demographic rates on climatic variables, it is well recognised that the response of an individual species to climatic change cannot be understood in isolation from the rest of the community (Davis et al. 1998, Pearson and Dawson 2003, Araújo and Luoto 2007, Gilman et al. 2010, Urban et al. 2012, 2016, Ockendon et al. 2014, Svenning et al. 2014, Ettinger and HilleRisLambers 2017, O’Brien et al. 2017, Legault et al. 2020). Although the relative balance of abiotic and biotic impacts on range edges may vary (Louthan et al. 2015, Paquette and Hargreaves 2021), rather than environment and competition acting as independent determinants of range limits, their combined effect is critical (Germain et al. 2018, Schultz et al. in press).

To date, the direct, systematic analysis of the effect of variability on extinction risk has been dominated by single-species studies (Bennie et al. 2013, Renton et al. 2014, Vasseur et al. 2014, Lawson et al. 2015, Bernhardt et al. 2018). Likewise, the majority of existing approaches to modelling the impact of climate change on multi-species distributions assume a smooth increase in the driving climate variable (Urban et al. 2012, Thompson and Gonzalez 2017, Alexander et al. 2018). However, temporal variability influences species dynamics, and hence response to climate change through multiple direct and indirect routes (Fig 1.)

**Figure 1.**
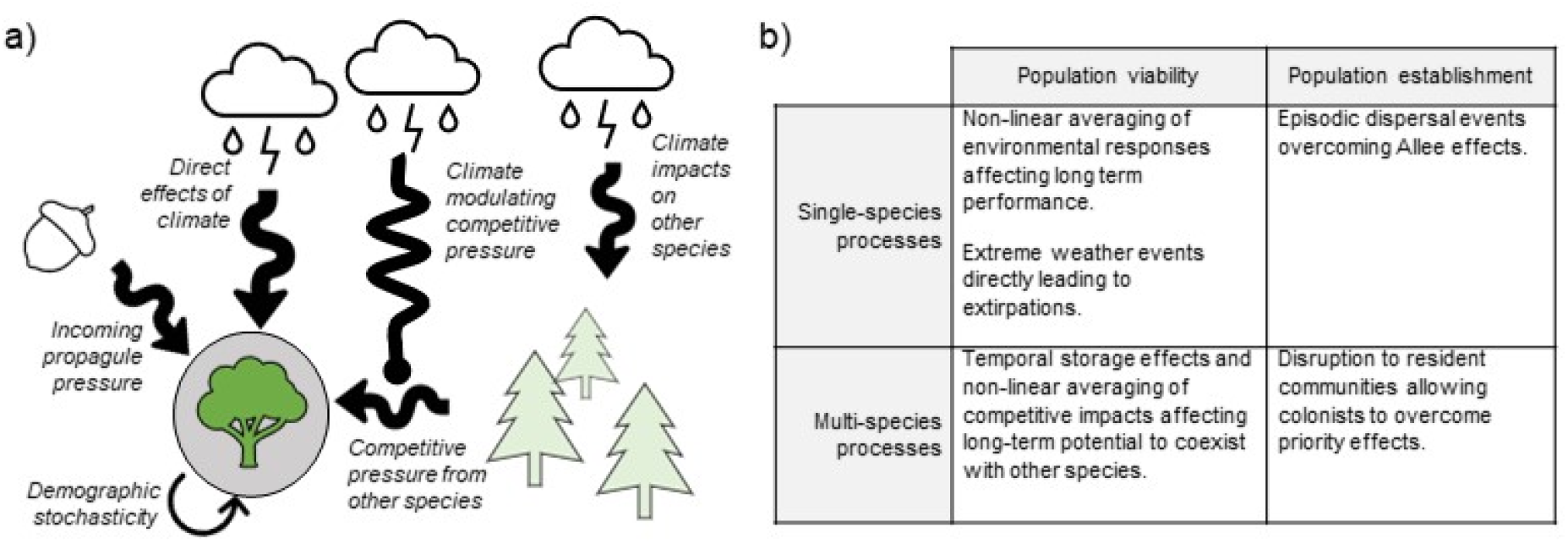
a) Schematic of principal routes by which variability influences a focal species at a site (circled). Climatic variability influences the focal population in three ways: directly influencing its reproductive rate, varying propagule pressure and through impacts of competitors (here represented as conifers). Overall competitive pressure can vary through fluctuating competitor numbers (which can be environmentally driven) and by varying modulation of the impact of competition exerted by the competitor. Variability generating processes internal to the focal population, such as demographic stochasticity, can interact with the externally driven variation. b) Categorisation of impacts of variability in determining species ranges and response to climate change discussed here. The mechanisms are categorised by organisational scale and whether they influence the ability for a species to persist at a site (its population viability) or the capacity for the species to establish new populations to shift their range.

We propose here that the current discussion around ecological responses to climate change could be missing a key element – the role of temporal variability in determining how interspecific interactions play out (Chesson and Huntly 1997, Vasseur and Fox 2007, Gudmundson et al. 2015, Fey and Vasseur 2016, Dee et al. 2020). Two distinct, but related, questions emerge: firstly, how does considering variability change our expectations for how communities will respond to shifts in mean conditions? And secondly, what impact would changes in environmental variability, for example interannual variance in temperature or rainfall (Swain et al. 2018, Chen et al. 2019), have on communities (Vázquez et al. 2017)?

Variability acts on each species in a community through numerous direct and indirect routes (Figure 1a). We structure our discussion by dividing the diversity of processes into those impacting the long-term viability of a population (Figure 1b), and processes affecting colonisation of new areas, drawing from ideas from invasion ecology. Using simple models, we then show how these processes can interact at all levels and how recent developments in techniques to partition the impact of variability can inform on the importance of different processes. We argue that the interrogation of parameterised models can help overcome the challenge of synthesizing insights from across ecological subfields.

We focus on populations near the edges of a species range, which constitute the front-line of climate change impacts -although we note that the geographic range edges are not always the most marginal habitats (Vilà-Cabrera et al. 2019, Oldfather et al. 2020). For vulnerable species with narrow climatic niches, their overall persistence will typically depend on their ability to advance their ranges. Species range edges are a unifying point of convergence for ecology (Holt and Keitt 2005) and as such the impact of climate change on ranges will require the synthesis of numerous strands of ecological thought (Urban et al. 2016). Populations can show greater sensitivity to climate variability towards the edge of their existing range, as observed for butterflies (Mills et al. 2017), tundra shrubs (Myers-Smith et al. 2015), game birds (Williams et al. 2003) and mangroves (Cavanaugh et al. 2018), although observational studies cannot easily distinguish whether these effects are direct environmental impacts or mediated through biotic interactions. The relative impact of biotic interactions is likely to vary across a species’ range, and can be particularly influential at range edges (Louthan et al. 2015, Early and Keith 2019, Bowler et al. 2020). We mostly focus on negative biotic interactions (particularly competition), but facilitation can have a key role in determining the range of some species (Bulleri et al. 2016), and the impact of facilitation is often higher in stressful locations such as towards range edges (Callaway et al. 2002, He et al. 2013).

We make a distinction when considering the impact of temporal variability between statistical descriptions of long-term patterns of fluctuations in climatic variables by measures such as variance and autocorrelation, and the impacts from individual discrete ‘extreme weather events’ defined either by their statistical unlikeliness or through the breaching of particular physiological thresholds (Smith 2011, Bailey and van de Pol 2016). The frequency of extreme events is expected to increase, due to both shifts in the underlying long term mean (which brings the threshold closer) and to increases in the underlying variance. The attribution of biological changes to specific anthropogenically-driven extreme events is possible, but demanding (Harris et al. 2020). Drawing a sharp line between long-term impacts of variance and discrete impacts is challenging, as individual extreme events are ultimately part of the ‘background’ variability observed over sufficient time. However, at the intermediate (multi-decadal) time scales of climate change concern, valuable insights can be gained from considering both aspects and synthesising the two is a key theoretical challenge.

### Variability and single-population growth rate

The direct impacts of climate variability on the viability of individual populations are widely appreciated (Lande 1993, Lawson et al. 2015), and so we only briefly review them here. Extensive analytical and experimental work demonstrates a long-term impact of a fluctuating climate on population growth rates (Ruel and Ayres 1999, Drake 2005, Melbourne and Hastings 2008, Thompson et al. 2013, Vasseur et al. 2014, Lawson et al. 2015). Average growth rates over the long term may be considerably different to population growth rates at average environmental conditions. Through non-linear averaging, the net impact of a variable climate on an individual species’ growth rate can be either positive or negative (Figure 2). The principal determinant of the direction of the effect is the curvature of the growth rate’s response to the relevant climate variable. However, higher-order properties such as temporal autocorrelation can also play a role (Petchey et al. 1997, Heino et al. 2000). Many populations near the edge of their range show greater population variability and climate sensitivity (Myers-Smith et al. 2015, Mills et al. 2017) and so may be expected to be particularly responsive to these effects.

**Figure 2.**
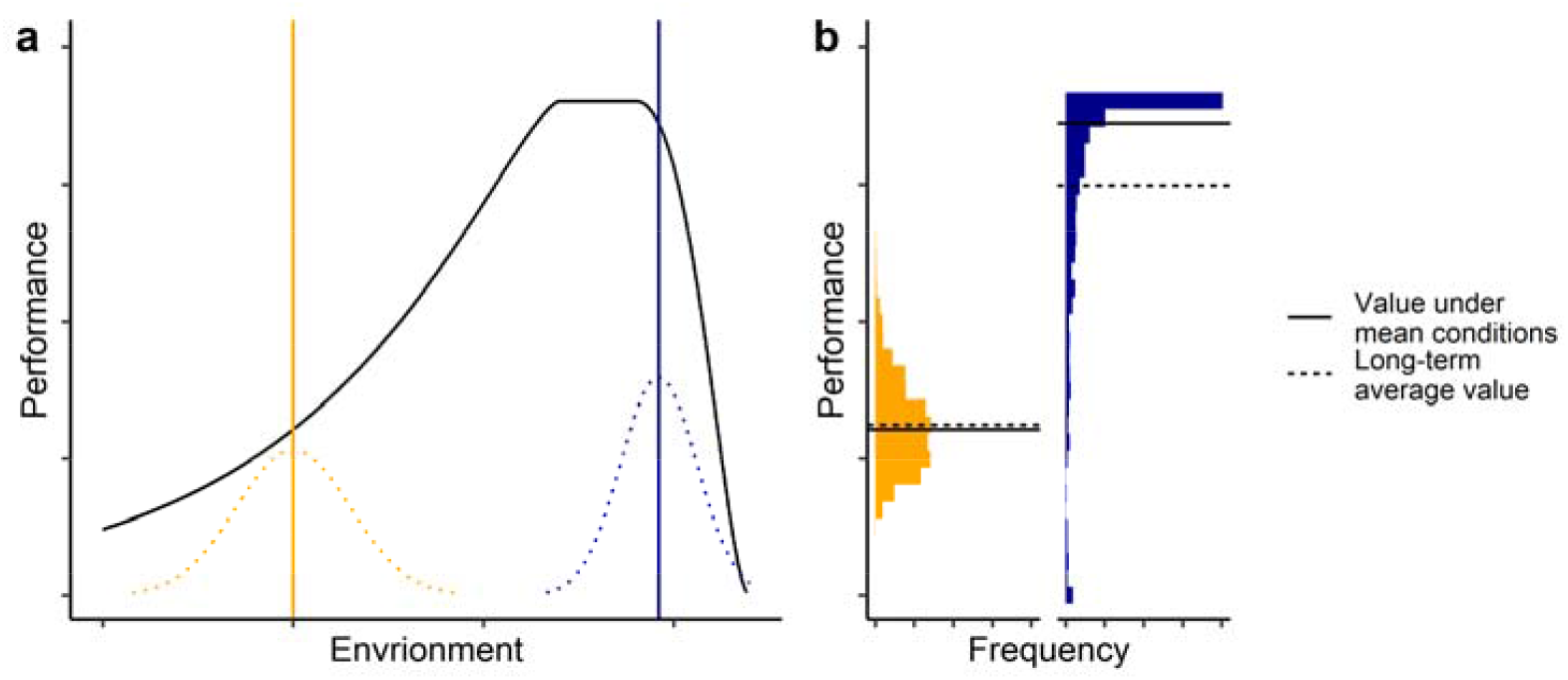
How the curvature of environmental performance curves (EPCs) affects average performance. a) The black line shows a classically shaped environmental performance curve where the key environmental variable is temperature, while the dashed lines show relative frequency of environmental conditions at two sites (yellow and blue). b) shows histograms of observed performances across 1000 random environmental draws at each site. At the yellow site, environmental variability is fairly large, but the curvature of the performance curve is relatively shallow. The long-term average value (dashed line) is therefore very similar to the value under mean conditions (solid line). At the blue site, although the variability is smaller, the local EPC curvature is much larger and downwards. The long-term value average value is considerably lower than the value under mean conditions. It also includes several instances that could be labelled extreme events, where the performance is very markedly below average.

Individual extreme events have most commonly been associated with population declines and heightened extinction risk, in particular when harsh climatic conditions push populations down to a level where they are vulnerable to extinction (Lande 1993, Boyce et al. 2006, Jongejans et al. 2010, Nadeau et al. 2017, Maxwell et al. 2019). Extreme events have been associated with increased extinction risk (Román-Palacios and Wiens 2020) and abrupt changes in community composition (Turner et al. 2020). Range contractions and community shifts have also been seen in many communities, including butterflies (De Palma et al. 2017), tropical fish (Lenanton et al. 2017), kelp (Smale and Wernberg 2013) and bumblebees (Soroye et al. 2020). Extreme events have led to extirpations from newly colonised areas, and it has been suggested that this may slow species responses to climate change (Nadeau et al. 2017). For example, extreme cold events have been associated with range retractions of invasive marine invertebrates (Canning-Clode et al. 2011) and fish (Rehage et al. 2016).

Taken together, both long-and short-term impacts of variability are more commonly viewed as having negative consequences for populations of conservation interest. However, as we shall show, when considering the wider community context in which species exist, this baseline assumption may need to be adjusted.

### Impacts of variability on long term coexistence

Where there is biotic control of species distributions, range limits become fundamentally a problem of coexistence (Shea and Chesson 2002, Usinowicz and Levine 2018). This framing unlocks for climate change research a long and rich history of work examining the influence of temporal environmental variability on coexistence (Levins 1979). The potential for temporal variability to enhance coexistence is well attested empirically (Adler et al. 2006, Tucker and Cadotte 2013, Tucker and Fukami 2014, Usinowicz et al. 2017, Hallett et al. 2019). Contrary to conclusions drawn in early and still influential literature (e.g. Hutchinson 1961), environmental fluctuations themselves are not sufficient to support coexistence of competing species (Chesson and Huntly 1997, Fox 2013) and can hinder as much as facilitate coexistence. The framework of modern coexistence theory (MCT, Chesson 2000) has been developed to robustly understand the impact of variability on long-term coexistence. However, this body of theoretical work examining the problem of species coexistence is only recently being applied to climate change in the context of spatially heterogeneous environments (Usinowicz and Levine 2018).

At its core, MCT defines and investigates coexistence in terms of the capacity for populations to expand from a low population density in the presence of competing species – summarised as the invasion criterion (Grainger et al. 2019b). Where all species are able to meet this criterion, they can each resist exclusion by the other species. In order to simplify the following discussion, we assume that only the ability of a particular focal species to persist at a site alongside one or more competitor species is in question. Through MCT, precise principles have been developed to identify how variability in the environment influences coexistence by quantifying the impact on the long-term average growth rate when the focal species is at low densities (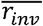, Chesson and Warner 1981, Chesson and Huntly 1997, Amarasekare et al. 2004, Snyder 2008).

MCT is grounded in analytical results which can be applied to a broad class of community models. This requires conceptually separating direct impacts of the environment on the focal population’s growth rate from the impacts exerted by competitors. This separation is delicate because competitors can affect the focal species directly, but also indirectly through shared resources - defined broadly to include physical resources such as nutrients, water or space, as well as through apparent competition mediated by natural enemy populations (Chesson and Kuang 2008). These routes of impact are the ‘limiting factors’ of MCT and can provide considerable analytical insight but also interpretational challenges (for a recent comprehensive review, see Barabás et al. 2018). However, as we shall show below, essential insights can be gained directly from a model that can describe how the growth rate of the population r depends on the environment (*E*) and the competitive impact (*C*), *r* = *f*(*E, C*), abstracting over the underlying mechanisms (Ellner et al. 2019). Where direct effects of environmental drivers and impacts of competitors contribute linearly and additively to the population growth of the focal species, any variability averages out in the long term. MCT can be used to describe how deviations from the linear additive base case lead to long-term effects of variability on population growth. These deviations can be understood in terms of two classes of impacts – temporal ‘storage effects’ and non-linearity of competitive effects.

Temporal storage effects arise when species differ in their response to the environment, such that the combined effects of the environment and competition allow benefits accrued in certain years to compensate for losses in other years (Chesson and Warner 1981). For this to affect long-term persistence two conditions must be met, in addition to species-specific environmental responses. Firstly, there must be an interaction between the direct impacts of the environmental conditions and the impacts of the competitor on the growth rate (mathematically this is non-additivity, where 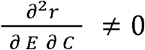. For example, a species may suffer proportionally less from competitors in a year of harsh environment if the population is buffered in some way such that the combined effect of a harsh environment and competitive pressure is capped (Fig 3ai). The classic mechanism that can lead to this ‘buffering’ is a resilient life stage such as a seedbank. Secondly, the impact of the environment on the focal species must be correlated through time with the competitive impacts on the focal species (i.e. E and C must co-vary, Figure 3aii). This is most directly generated by specialisation of species to different environmental conditions. For example, if the focal species performs comparatively better in cool temperatures, in a cold year it would respond well to the environment and the competitor species may impose less competitive pressure. Alternatively, this covariance can be generated through temporal autocorrelation in environmental conditions (Schreiber 2021).

**Figure 3.**
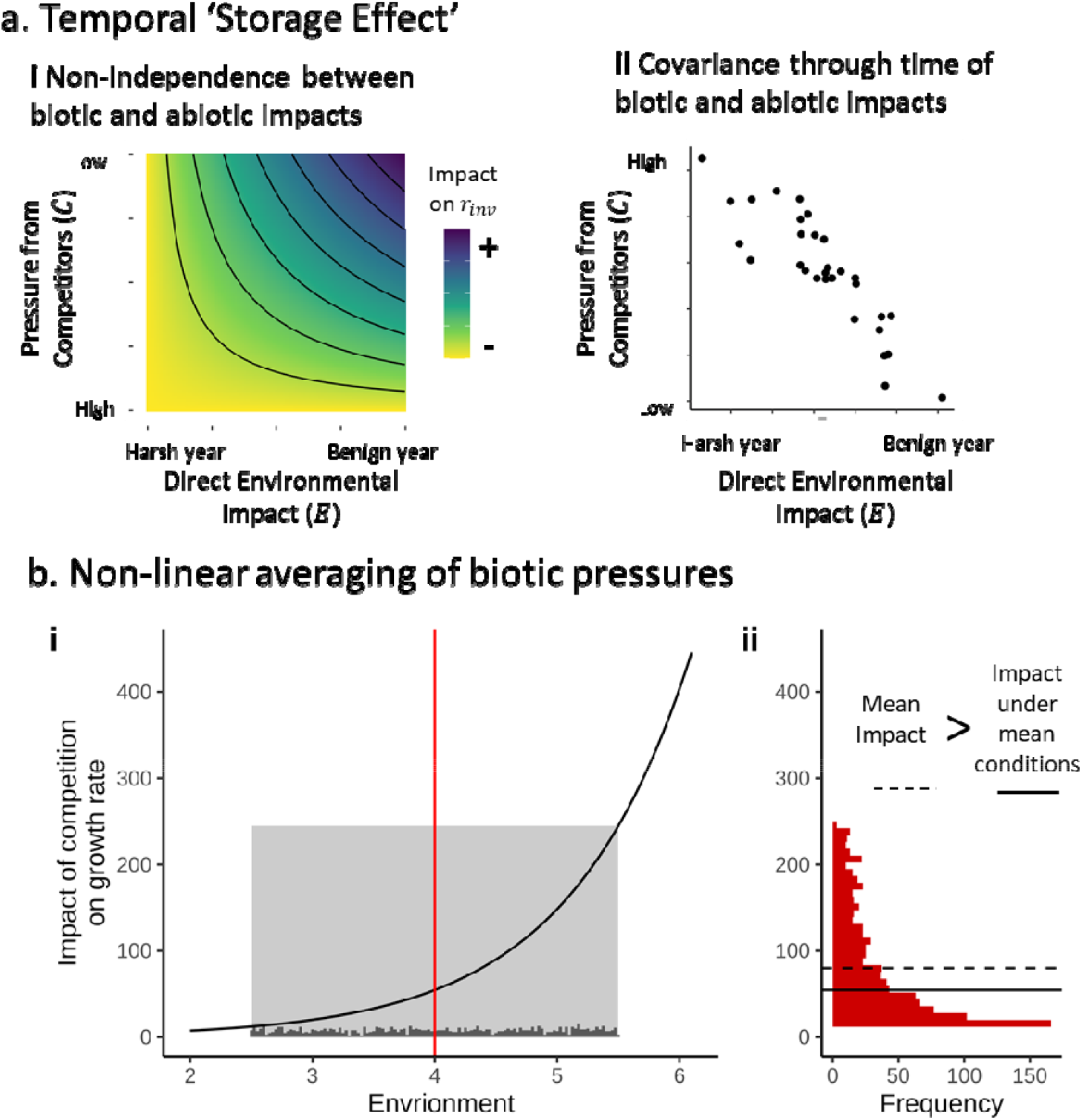
Illustration of the conditions required for the two multi-species mechanisms by which environmental variability can impact persistence. a) Temporal storage effects describe how species that differ in environmental dependence are able to divide niche opportunities afforded by a variable climate between them. For this to influence the long-term average population growth rate, there are two conditions: i) the impacts of competitors and the environment must interact in determining growth rate, and ii) the impact of competition and direct effects of the environment must correlate through time. Here each dot represents a year and pressure from the competitor tends to be lower in a benign year for the focal species. This could derive from the two species prospering in different conditions. b) Non-linear averaging of the impact of biotic pressures can result in an overall impact on average population growth rate that differs from the impact under mean conditions. Here we show a case of environmentally determined impact of competition. The determinants of the net effects of non-linearities of biotic impacts have similar drivers to the single-species mechanisms described in Fig.2, depending on both the pattern of environmental variability and the curvature of species responses.

In the classic case with buffered (‘subadditive’) population growth, the more negative this covariance, the greater the beneficial effect to the focal population. However, it is worth noting that temporal storage effects can be reversed if the biotic and abiotic impacts on the focal species growth rate are ‘superadditive’, i.e. the adverse effects of competition are proportionally greater in a harsh year (e.g. Holt & Chesson 2014). In the context of climate change, there is a risk that the current patterns of covariation in species’ responses to the environment could change. For example, if climate change results in more frequent universally ‘bad’ (or equally, universally ‘good’) periods, instead of a back-and-forth of alternate species being favoured, overall covariance in species responses could become more positive and further undermine coexistence.

The second mechanism arises directly from fluctuations in the impact of other species on the growth rate of the focal species. Non-linear averaging of varying biotic impacts on the focal species’ growth can affect 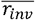, analogous to the non-linear averaging of abiotic environmental fluctuations on population growth rates described in the section above (Fig 2), and with matching consequences for shifts in climatic variability patterns (Fig 3b). These fluctuations in the biotic pressure can be driven by changes in the abundance of competitor species, or via varying per-capita intensity of the competition exerted by other species through fluctuations in shared resources. When examining coexistence, this is the mechanism of ‘relative nonlinearity’ that describes how species can differentiate themselves through their capacity to take advantage of variable environments - ‘slow-and-steady’ versus ‘boom-or-bust’ dynamics (Armstrong and Mcgehee 1980). Notably, in contrast to temporal storage effects, this does not directly rely on correlations in species response to the environment and can also derive from other fluctuation generating mechanisms.

The foundation for identifying the relative strengths of these processes in a real system is the construction of a simple parameterised model. With that in hand, approaches such as that proposed by Ellner *et al*. (2019) can be used to partition — into contributions of different single-species and multi-species aspects of variability, without the need for complex analytical work and overcoming limitations incurred by approximations made in the analytic theory (Hallett et al. 2019, Zepeda and Martorell 2019, Shoemaker et al. 2020b). We describe this approach in Figure 4 and in SI 2. In a recent example, Armitage and Jones (2020) used a model of competition between two species of duckweed to show that the inferior competitor’s poleward range limit is better predicted when taking into account the impact of temporal fluctuations. Using a partitioning approach, they found this was dominated by nonlinearity in direct temperature responses, with a smaller contribution of non-linearities in competition and minimal impact of temporal storage effects attributable to the positively correlated species responses.

**Figure 4.**
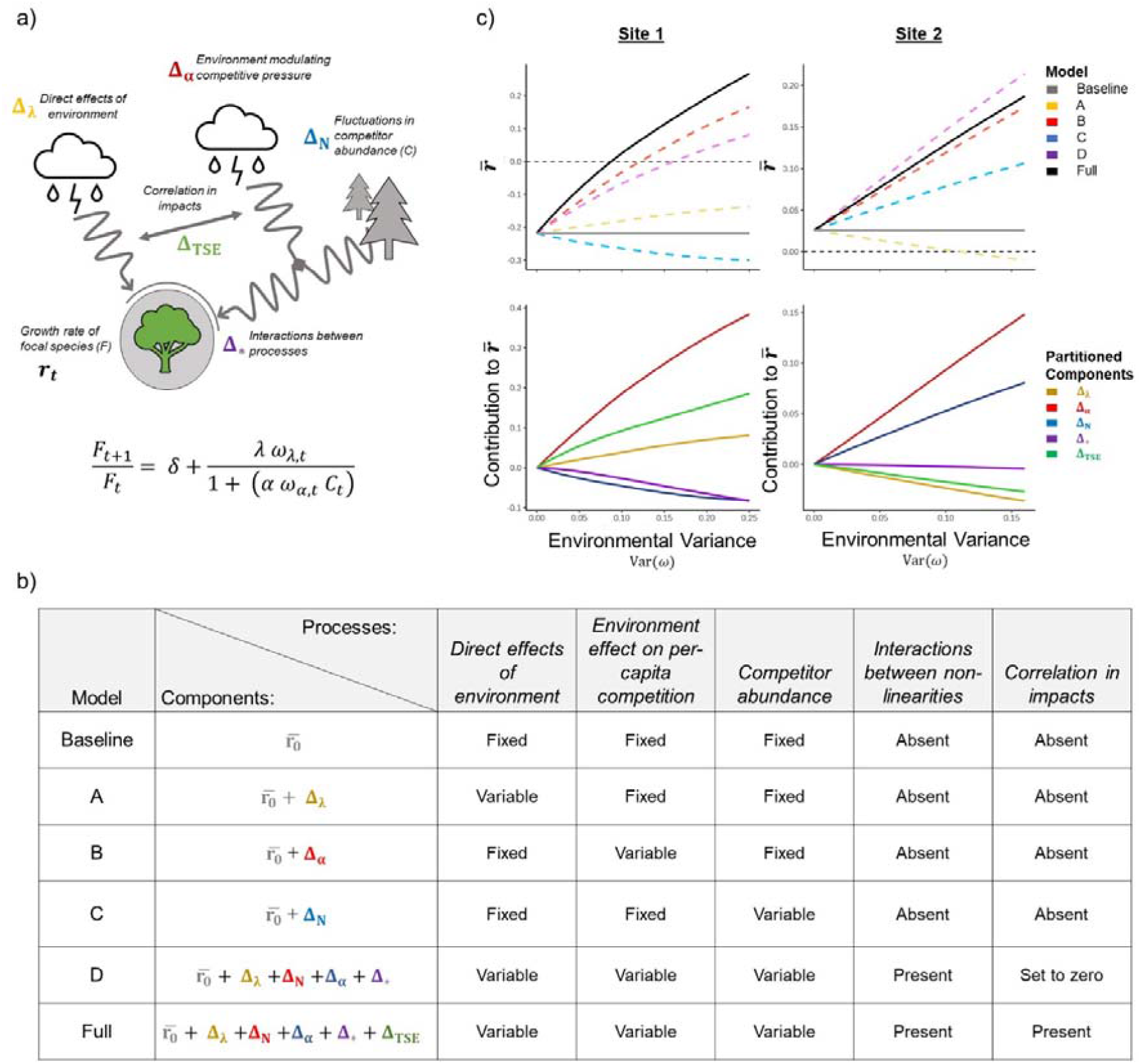
Summary of the steps required to investigate impact of temporal variability through the partitioning method of Ellner et al. (2019) and two examples. Full details are given in SI 2. a) Firstly, establish the processes involved and build a model of the population dynamics to calculate the long-term growth rate 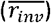. In our model, rate of population change when rare at each time point is given by Eq (1), where δ = carryover from the previous generation, λ = fecundity in the absence of competitors, α = per-capita competitive coefficient. Since we assume the focal species is rare it includes no intra-specific competition. The impact of a variable climate on competition and intrinsic growth rate at each time step is determined by the two ω terms, which are correlated at each time step t. The population density of the competitor (*C*_*t*_) also fluctuates through time.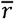 is found from the mean of log (*F*_*t*+1_/*F*_*t*_) over a sufficiently large number of trials. b) Secondly, systematically alter the inputs of the growth rate model by sequentially fixing different fluctuating terms to their average values or removing correlations between variable components. Choices of which aspects to fix, and in which combination will allow different partitioning. Here we partition the difference between the growth rate under fully fixed conditions 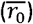 and under the ‘observed’ variable conditions 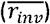 with five Δ-partitions: 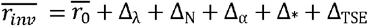, corresponding to the processes identified in a). Differences between the model variants allow the identification of the relative influence of each partition. c) Examples of how the partitioning approach can illustrate how variability can have multiple counteracting consequences for a species’ ability to persist. We use two example parameterisations of the same underlying model, Site 1 (left) and Site 2 (right). At Site 1 without variability, 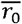 is negative, and the focal species would not be expected to persist. The full model (black line) suggests that with a climate variability above 0.08 persistence is possible. Inspection of the partitions (below) shows that the two largest positive impacts are non-linearities in the impact of competition (Δ_α_) and temporal storage effects (Δ_TSE_). Opposing this, variability in the abundance of competitors (Δ_N_) is detrimental to the focal species. The impact of variability directly on the growth rate of the focal species (Δ_λ_) is here relatively small, and insufficient to lead to persistence in the range of variabilities examined. The interaction term between the processes (Δ_*_) is negative but small. Note that at low levels of variability Δ_*_is slow to increase, showing that the other partitions identify the main pathways of low variability levels. By contrast, at Site 2, variability in the growth rate of the focal species has a negative impact (Δ_λ_), and if it was the only process considered might be expected to prevent persistence at the site. There is also a negative temporal storage effect (Δ_TSE_), deriving from a positive correlation between competitive impact and intrinsic growth rate in the underlying model. However, these two effects are more than counteracted by strong positive contributions from non-linearities in both competitive effect (Δ_α_) and abundance of competitors(Δ_N_). In this case the role of interactions between non-linearities (Δ_*_) remains small.

Earlier theoretical work placed greater emphasis on temporal storage effects, but in the small number of empirical cases where the relative impact of the two effects on coexistence has been directly compared, relative nonlinearity was found to have comparable, or greater, impact than the more widely appreciated storage effects (Letten et al. 2018, Hallett et al. 2019, Zepeda and Martorell 2019). Although to date the number of examples is currently small, it is clear that climate-driven shifts in variability patterns could play a role in determining coexistence between competitors and range limits in the future. It is also clear that the impact of long-term trends in average values need to be understood in in terms of underlying variability which may help or hinder coexistence. While we have focussed here on competitive systems, consumer-resource systems can be analysed in parallel ways (Shoemaker et al. 2020b, Dee et al. 2020). However, effects identified in simple communities may not necessarily directly translate to more complex systems (Barabás et al. 2018, Song et al. 2019). Species respond to different parts of the environment - with a greater diversity of species, it is quite possible that species-level fluctuations in competition may average out at the community level. For example, Clark *et al*. (2010) found that different tree species responded to different aspects of the overall environmental fluctuations. This may suggest that these mechanisms may be strongest where a limited pool of species are involved in constraining the range of the focal species.

### Community impacts of discrete events

A species’ range expansion in response to climate change is effectively a series of invasions into new communities (Wallingford et al. 2020). From this perspective, the significant body of work investigating variability within invasion biology, going back to Elton (1958) and beyond, can offer useful insights, important distinctions notwithstanding (Urban 2020). The ability of a species to persist at a site is only one half of the picture – a species must first arrive and establish itself. The spread of a species into new areas can be slowed or even prevented by disadvantages that potential invaders face. For instance, positive density dependence at low population densities (Allee effects, Courchamp et al. 1999, Kramer et al. 2018) can cause leading range edges to appear ‘pinned’ in place (Keitt et al. 2001) and slow the rate of invasion into newly suitable environments (Taylor and Hastings 2005).

Environmental variability can play a role in shifting a community from one state to another by allowing species to overcome the challenges of Allee effects through intermittent boosts in performance (Dennis 2002). Discrete extreme weather events can have marked influence on the trajectory of species responses to climate change, but pose considerable challenges to investigation and prediction (Bailey and van de Pol 2016). Although direct evidence is challenging to find (see later sections), extreme climatic events have been associated with the arrival into marine communities of species previously found in warmer areas (Wernberg et al. 2013). Dispersal is intrinsically episodic (e.g. Kennedy et al. 2020) and short term spikes in the number of incoming colonist propagules may help increase establishment compared to constant dispersal rates by overcoming thresholds induced by Allee effects (Drake and Lodge 2006, Carr et al. 2019).

At the community level, biotic resistance from the resident species can slow or prevent a colonist tracking its climatic niche (Urban et al. 2012, Legault et al. 2020). Whether this resistance is considered a hindrance or beneficial will depend on the conservation status and impact of the colonist and resident species concerned. Over longer time scales, a history of disturbance can shape a community’s biotic resistance through selective assembly (Miller et al. 2021) or through specific adaptations to the local conditions, which may include levels of variability (Urban and De Meester 2009), although temporal variability in climate could also preclude such local adaptation from occurring (Bridle et al. 2010).

Even without local adaptation, priority effects can give residents considerable advantages compared to potential invaders. Where priority effects are strong, an invading species can colonise only if either the density of the resident is brought down from equilibrium, or the invader is otherwise able to reach sufficiently high densities to exert significant competitive pressure on the resident. Individual disturbance events can temporarily break down blocking effects (Davis et al. 2000, Melbourne et al. 2007, Diez et al. 2012), for example in grasslands (Pinto and Ortega 2016) and over the longer term there is an expectation that in disturbed environments there are more unused available resources for invaders to take advantage of (Davis et al. 2000, Diez et al. 2012). Tucker and Fukami (2014) showed experimentally that temperature variability can allow priority effects to be overcome in a nectar-yeast system. Ecological theory can play a key role in identifying cases where individual key events could precipitate the establishment of a climate refugee species. The core results of MCT can be applied to invasions (Shea and Chesson 2002, MacDougall et al. 2009) and are a useful guide to identifying where priority effects are impactful (Ke and Letten 2018, Uricchio et al. 2019, Grainger et al. 2019a) particularly where the growth rate of a colonising species can also be affected by variability (Clark and Johnston 2011).

### Tackling interactions between influences of variability at multiple scales

When considering whole communities, the diversity of possible impacts on a focal species due to variability is considerably larger than in the single-species case. The previous three sections demonstrated the breadth of direct and indirect ecological impacts that variability can have on how species will respond to climate change. Faced with such a diverse set of processes, reconciling the assorted influences of variability and determining how they interact is central to determining their influence in practice. There are fundamentally different scales and mechanisms at work, but bottom-up mechanistic modelling can illustrate the key interactions at play. Identifying and (equally importantly) ruling out for practical purposes, interactions between stressors is crucial to meaningful conservation interventions (Côté et al. 2016).

There have been frequent calls to improve the representation of communities in ecosystem-change models (Gilman et al. 2010, Angert et al. 2013, Urban et al. 2013). This poses a considerable challenge, as directional climate change models are fundamentally non-stationary, requiring fundamental extensions beyond the study of equilibria analysis central to classical ecological theory (Chesson 2017). Synthesising the aggregate impact of variability will require an expansion in the scope of models currently used (Felton and Smith 2017).

To this end, a number of modelling studies have explored the interface between local variability-mediated coexistence and extinction risk (Adler and Drake 2008, Gravel et al. 2011, Danino et al. 2018, Schreiber et al. 2019, Pande et al. 2019, Dean and Shnerb 2020). Populations at low densities may be expected to benefit the most from variability-mediated coexistence mechanisms, but a low population size is also risky if a single bad year could extirpate the population. In these models, the relative strengths of stochastic extinction risk and competitive stabilization change across a gradient of environmental variability. In the largest empirical analysis of this balance to date, Fung *et al*. (2020) used forest plot data to quantify how variability leads to temporal niche partitioning and extinction risk and found that the balance was uneven but more frequently detrimental to coexistence.

A useful way of framing complex climate change responses into a single unified measure of impact is through establishment and extinction lags - differences between climate change and species range responses (Alexander et al. 2018). The core issues can be demonstrated in relatively simple simulation models constructed to capture multiple processes and forms of variability simultaneously. As an example, in SI 3 we describe a simple discrete-time model of the population dynamics of a resident species and a climate migrant that are in competition with each other and subject to various types of variability - in baseline performance, in propagule pressure, and in demographic stochasticity. In Figure 6 we demonstrate how the average time for the colonising species to become fully established responds to an aspect of variability can be dependent on complex interactions with other details of the model, with the effect that the response expected from a change in variability due to one mechanism could be countered or even reversed in conjunction with other processes. Given the multitude of theoretically and empirically identified effects of climate variability on colonisation success under climate change, there is a need to develop and investigate such models to understand when interactions between these effects are likely to be influential. Comparison between asymptotic predictions made under frameworks such as MCT and predictions made under stochastic frameworks allows the identification of the impact of discrete variability and direct testing of the assumptions of long-term averages of MCT.

**Figure 5.**
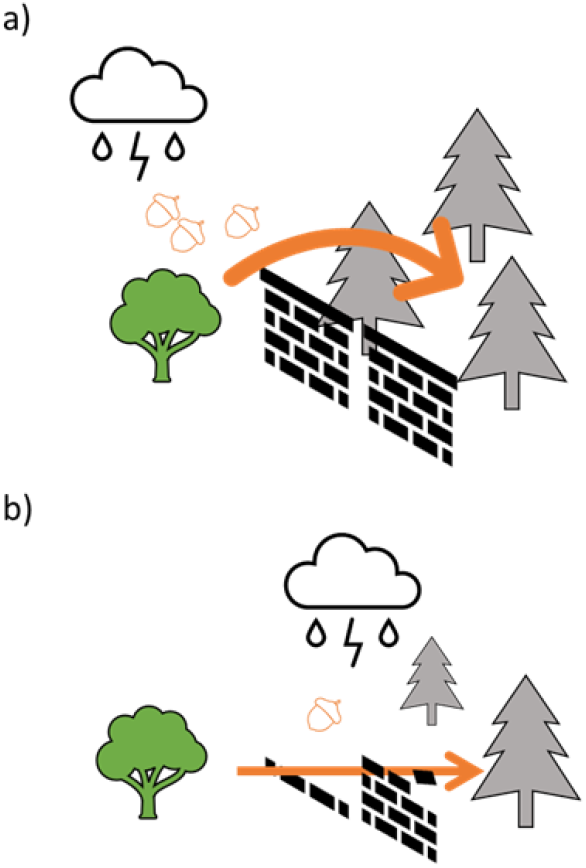
Illustration of two ways in which climatic variability can overcome barriers to range shifts. a) through episodic high-dispersal events, and b), by disrupting existing communities and reducing biotic resistance.

**Figure 6.**
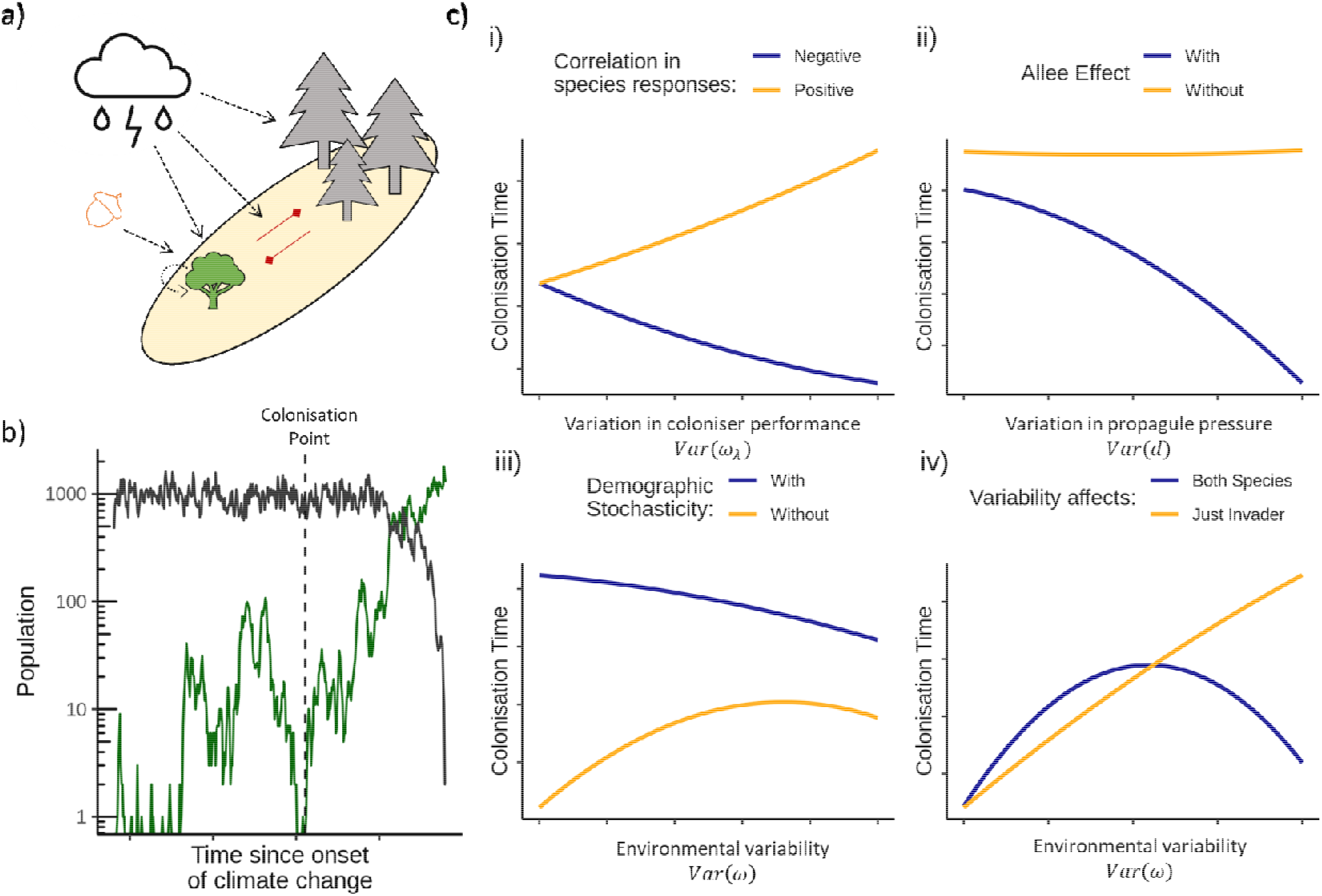
Demonstration of interactions between impacts of environmental fluctuations on climate-driven colonisation in a simple two-species competition model. a) The core model is an extension of that discussed in Figure 4 and is detailed in SI 3. Initially the site is dominated by a resident species and subject to immigration pressure from a potential coloniser. Over time, the environment becomes increasingly favourable for the coloniser and eventually it will displace the resident. We introduce dynamical features or additional sources of variability and observe how average colonisation times respond to changes in certain aspects of variability. Here, we measure the time point from the onset of climate change when the coloniser is permanently established (b). c) Four examples of interaction between model processes, shown with smooth lines through 500 trials: i) as variability in the coloniser’s performance rises, colonisation occurs more rapidly with increased variability where there is a negative correlation (blue) between the performance of the resident and the coloniser, as temporal storage effects aid establishment. However, where the competitor responses are positively correlated (yellow) colonisation is slowed by variability, ii) variability in incoming propagule pressure accelerates the colonisation only where there are strong Allee effects to overcome, iii) a colonisation delay with environmental variability can be reversed where there is demographic stochasticity that can lead to extinction at low densities, iv) the beneficial effect of variability for the coloniser can be reversed when it affects both species.

At a landscape scale, regional processes can influence responses to local directional change (Anderson et al. 2009, Kahilainen et al. 2018, Ryser et al. 2021) particularly if frequent local extirpations towards range edges drive metapopulation dynamics. Building on small and focused models, larger, highly generalised and spatially-explicit metacommunity models (O’Sullivan et al. 2019, Thompson et al. 2020) can provide insight into potential drivers of community change that emerge from combining processes at multiple scales (Usinowicz and Levine 2018, Chase et al. 2020). However, interpreting such models to assess the impact of variability poses distinct challenges, beyond parameterisation. It is rarely possible to directly control multiple aspects of variability simultaneously, even in an artificial model. Temporal variability is inherently multi- facetted and additional qualities beyond direct variance can have significant impacts, e.g. autocorrelation (Levine and Rees 2004). To take one illustrative example, the historical level of variability a community (real or *in silico*) experienced during its assembly contributes to the capacity of the community to respond to future changes, whether that is through direct adaptation of the species in a community to local levels of variability or by the extant species having passed through a previous extinction filter during historical extreme events (Janzen 1967, Nadeau et al. 2017, Medeiros et al. 2020, Miller et al. 2021).

## Identifying processes in the real world

The next frontier is directly assessing the magnitude of these effects in real systems which will require empirical work. Possible driving features of climate sensitivity such as latitudinal patterns appear to have very mixed effects (Louthan et al. 2021). Determination of the relative contributions of dispersal, interspecific interactions and environmental dependence has been identified as the key challenge to understanding the dynamics of whole communities (Leibold et al. 2021). There is evidence that biotic resistance to invasive species is widespread, but the global contribution of biotic resistance to climate refugee species is challenging to measure (Levine and Rees 2004, Alexander et al. 2015, 2016, Louthan et al. 2015, Godsoe et al. 2017, 2018, Beaury et al. 2020). Understanding which aspects of a variable environment are most influential will be key to building models of minimal necessary complexity. Correspondingly, choosing appropriate time resolution is critical - where competing species have different life-cycle periods, fluctuations on different frequencies (such as seasonal migration or environmental changes) can have emergent impacts (Hutchison et al. 2020).

Direct observations demonstrate that species are on the move, but consistent patterns are difficult to determine and influenced by concurrent land use changes (Lenoir et al. 2020). The observed rate of change in species ranges is highly variable, with many species shifting their ranges considerably faster or slower than the climate velocity and ultimately dependent on availability of habitat (Platts et al. 2019). Competitive exclusion at large spatial scales is often very slow (Yackulic 2017), while extirpation by extreme events can be rapid, but not necessarily permanent. Any coupling between species ranges and particular climatic events can be highly idiosyncratic, with multi-year effects of weather events (Harley and Paine 2009). Coupled with the challenge of accurately identifying the pace and cause of range shifts (Bates et al. 2015, Taheri et al. 2021), this makes directly discerning a signal of variability in movement rates an imposing task.

Despite this, direct observations of variability in natural populations can highlight how species respond differently to environmental variability (Palmer et al. 2017, Le Coeur et al. 2021). Evidence from global satellite data shows that sensitivity to climate variability is itself variable across the globe (Seddon et al. 2016). However, to determine the impact in terms of long-term coexistence, model parameterisation of some sort is required (e.g. Fung et al. 2020, Usinowicz and Levine 2021). Species traits hold some promise to identify likely temporal coexistence mechanisms (Adler et al. 2013). Life history traits have been found to relate to sensitivity to climate anomalies in herbaceous perennials (Compagnoni et al. 2021), birds (Cohen et al. 2020), and amphibians (Cayuela et al. 2017), but much work remains to be done in this area.

Mesocosm experiments with manipulation of variability can be illuminating – for example, Zander et al. (2017) showed that lower trophic levels of a microbial food web were more strongly affected by variability than top level consumers. However, such an approach is fundamentally limited since variability can be manipulated in many alternative valid dimensions unless it is tied directly to expected climate regimes (Thompson et al. 2013). Behavioural adaptation and the role of microclimates pose further challenges to the interpretation of mesocosm work - the realised variability of environmental variables relevant to species may differ from that measured by weather stations (Bladon et al. 2020).

Alongside the highly generalised ‘strategic’ models demonstrated in the previous section, multiple impacts of variability need to be tested for in focussed ‘tactical’ case studies of marginal populations in order to build a picture of the real-world prevalence of these processes. Progress will require not just more data, but a connected approach to synthesising the multiple impacts of variability, which in turn requires a reliable model of community dynamics that can incorporate variable conditions. At the core will be robust models of species performance and competitive impact under different environmental conditions. This is no easy task – even in two-species systems with a single environmental variable this requires fitting a multidimensional response surface. Guidelines for minimal experimental designs to assess competition have been established (Hart et al. 2018) and extensions to alternative environments can be quite direct. Although experiments require significant investment, it is not out of reach. For example, Hallett et al. (2019) successfully parameterised a model of grass-forb competition under variable rainfall using four environmental treatments crossed with ten species density treatments. In complex communities, modelling variation in overall competitive pressure on a focal species may be a necessary abstraction. Given that species respond to multiple environmental variables (Clark et al. 2010, Tingley et al. 2012) there are fundamental limits to the resolution such models can aim to achieve. Nonetheless, we believe that an increased emphasis on parametrised models as a core goal for empirical research would bring marked benefits. Detailed predictions for individual communities will need to confront measures of environmental performance with observations of current ranges to best estimate future trajectories (Armitage and Jones 2020, Twiname et al. 2020). Observations of communities along climate gradients can provide particularly valuable tests of models designed to investigate climate change (Alexander et al. 2016, Tylianakis and Morris 2017).

With a sufficiently supported model in hand, partitioning of the various impacts of variability on coexistence can be highly informative (Figure 4, Ellner et al. 2019, Shoemaker et al. 2020b). The method can be extended to the large scales relevant to climate-change responses (Armitage and Jones 2020). Such a simulation-based partitioning approach could also be applied with other response variables, for example colonisation or extirpation lags, or extended to include additional variability terms. Understanding which aspects of variability are most influential can sharpen the focus of investigations, reducing overall problem complexity and concentrate potential mitigation efforts on the most critical fluctuations.

In support of this, a key line of future theoretical enquiry will be determining the minimum data requirements to understand the impact of variability. It is not yet known how sensitive partitioning of variability effects is to model misspecification. The higher-level properties of environmental performance curves, such as their curvature, are considerably harder to estimate than first order properties such as thermal optima. Empirical estimates for key parameters can be confounded with each other, with consequences for reliable estimation of species coexistence (Terry et al. 2021). The implications of parameter uncertainty need to be explicitly acknowledged and better understood. Since partitioning is algorithmic, it is possible to propagate uncertainties in the underlying model through to uncertainty in the impact of variability on the response of interest.

## Conclusion

In Table 1 we summarise the multitude of ecological routes by which underlying temporal variability could influence how a species will fare under climate change. It is not currently clear whether the difference in emphasis of the impact of variability in different ecological subfields represents a lack of communication, publication bias, or if the relative neglect of the ‘positive’ aspects of variability within global change biology is because they do not leave a widespread strong imprint on real-world dynamics. It would be risky to assume that the impacts of variability are already ‘baked-in’ to current observed species ranges, and so captured by existing distribution models. We reiterate that there will also be evolutionary processes to consider - interactions between variability and adaptation with positive and negative consequences for range shifts has been subject of extensive recent reviews elsewhere (Vázquez et al. 2017, Nadeau and Urban 2019, Thompson and Fronhofer 2019, Miller et al. 2020, Coleman and Wernberg 2020, Lyberger et al. 2021).

**Table 1.** Summary of the consequences climate variability can have on species responses to climate change. Influential interspecific interactions are necessary for the multi-species process to be impactful. Particularly in the multi-species processes, more research is needed to determine their prevalence and influence in real systems.

| Process | Key determinant of impact | Possible impact of considering variability in the response to climate change |
| --- | --- | --- |
| *Single Species* |  |  |
| Non-linear averaging of environmental responses | Curvature of environmental performance curve | Long-term population viability could differ from viability under average conditions |
| Extreme weather-related population impacts | Population sensitivity and recovery rates | Generally considered detrimental to population persistence |
| Episodic dispersal | Dispersal limitation of range expansion<br>Allee Effects | Establishment into new areas may be aided |
| *Multi-Species* |  |  |
| Temporal storage effects | Covariance between environmental and competitive impacts on growth rate<br>Non-additivity of environmental and competitive impacts | Consistency of species' environmental responses could be influential in determining persistence |
| Non-linearity of competition/competitive impact | Curvature of competitive impact on growth rate | Long-term population viability could differ from viability under average conditions |
| Disruptions overcoming priority effects | Strength of biotic resistance due to priority effects<br>Disturbances reducing biotic resistance | Biotic resistance could be overcome more suddenly than expected |

At this point in time, we simply do not know whether current assumptions of the impact of variability based on single-population analyses are systematically over or underestimating risks at the community level. What we do know is that climate change will present species with a bumpy and obstacle-filled uphill ride, not a smooth escalator. No single simple theory can predict the effects of climate variability – but, as we have shown, this does not prevent useful insights emerging from the synthesis of multiple subfields of ecology. Future theoretical work should aim to identify not just possibilities, but the circumstances in which these processes are impactful. As with most areas of ecology, both complex and simple verbal and mathematical models have their parts to play. By understanding the linkages between these models, detailed insights can be gained without losing sight of the whole. More examples of quantification of the impact of variability in real communities are needed - it is our belief that the simple modelling frameworks discussed here can meet this need. Building strong bridges between climate change ecology and coexistence theory has never been more possible, or more necessary.

## Authorship

JCDT wrote the manuscript and built the models. All authors contributed significantly to the editing and manuscript development.

## Data Sharing and Data Accessibility

Code to generate all results is publicly available at https://github.com/jcdterry/ClimateVar_BioticInts and should the manuscript be accepted will be permanently archived on Zenodo. The paper contains no new datasets.

## References

Adler, P. B. and Drake, J. M. 2008. Environmental variation, stochastic extinction, and competitive coexistence. - American Naturalist 172: E186–E195.

Adler, P. B. et al. 2006. Climate variability has a stabilizing effect on the coexistence of prairie grasses. - Proceedings of the National Academy of Sciences 103: 12793–12798.

Adler, P. B. et al. 2013. Trait-based tests of coexistence mechanisms. - Ecology Letters 16: 1294–1306.

Alexander, J. M. et al. 2015. Novel competitors shape species’ responses to climate change. - Nature 525: 515–518.

Alexander, J. M. et al. 2016. When Climate Reshuffles Competitors: A Call for Experimental Macroecology. - Trends in Ecology & Evolution 31: 831–841.

Alexander, J. M. et al. 2018. Lags in the response of mountain plant communities to climate change. - Global Change Biology 24: 563–579.

Amarasekare, P. et al. 2004. Mechanisms of coexistence in competitive metacommunities. - American Naturalist 164: 310–326.

Anderson, B. j et al. 2009. Dynamics of range margins for metapopulations under climate change. - Proceedings of the Royal Society B: Biological Sciences 276: 1415–1420.

Angert, A. L. et al. 2013. Climate change and species interactions: ways forward. - Annals of the New York Academy of Sciences 1297: 1–7.

Araújo, M. B. and Luoto, M. 2007. The importance of biotic interactions for modelling species distributions under climate change. - Global Ecology and Biogeography 16: 743–753.

Armitage, D. W. and Jones, S. E. 2020. Coexistence barriers confine the poleward range of a globally distributed plant. - Ecology Letters 23: 1838–1848.

Armstrong, R. A. and Mcgehee, R. 1980. Competitive Exclusion. - The American Naturalist 115: 151–170.

Bailey, L. D. and van de Pol, M. 2016. Tackling extremes: Challenges for ecological and evolutionary research on extreme climatic events. - Journal of Animal Ecology 85: 85–96.

Barabás, G. et al. 2018. Chesson’s coexistence theory. - Ecological Monographs 88: 277–303.

Bates, A. E. et al. 2015. Distinguishing geographical range shifts from artefacts of detectability and sampling effort. - Diversity and Distributions 21: 13–22.

Beaury, E. M. et al. 2020. Biotic resistance to invasion is ubiquitous across ecosystems of the United States. - Ecology Letters 23: 476–482.

Bennie, J. et al. 2013. Range expansion through fragmented landscapes under a variable climate. - Ecology Letters 16: 921–929.

Bernhardt, J. R. et al. 2018. Nonlinear averaging of thermal experience predicts population growth rates in a thermally variable environment. - Proceedings of the Royal Society B: Biological Sciences 285: 20181076.

Bladon, A. J. et al. 2020. How butterflies keep their cool: Physical and ecological traits influence thermoregulatory ability and population trends. - Journal of Animal Ecology 89: 2440–2450.

Boettiger, C. 2018. From noise to knowledge: how randomness generates novel phenomena and reveals information. - Ecology Letters 21: 1255–1267.

Bowler, D. E. et al. 2020. Impacts of predator-mediated interactions along a climatic gradient on the population dynamics of an alpine bird. - Proceedings of the Royal Society B: Biological Sciences 287: 20202653.

Boyce, M. S. et al. 2006. Demography in an increasingly variable world. - Trends in Ecology and Evolution 21: 141–148.

Bridle, J. R. et al. 2010. Why is adaptation prevented at ecological margins? New insights from individual-based simulations. - Ecology Letters 13: 485–494.

Bulleri, F. et al. 2016. Facilitation and the niche: implications for coexistence, range shifts and ecosystem functioning. - Functional Ecology 30: 70–78.

Callaway, R. M. et al. 2002. Positive interactions among alpine plants increase with stress. - Nature 417: 844–848.

Canning-Clode, J. et al. 2011. ‘Caribbean Creep’ Chills Out: Climate Change and Marine Invasive Species (M Peck, Ed.). - PLoS ONE 6: e29657.

Carr, A. N. et al. 2019. Long-term propagule pressure overwhelms initial community determination of invader success. - Ecosphere 10: e02826.

Cavanaugh, K. C. et al. 2018. Sensitivity of mangrove range limits to climate variability. - Global Ecology and Biogeography 27: 925–935.

Cayuela, H. et al. 2017. Life history tactics shape amphibians’ demographic responses to the North Atlantic Oscillation. - Global Change Biology 23: 4620–4638.

Chase, J. M. et al. 2020. Biodiversity conservation through the lens of metacommunity ecology. - Annals of the New York Academy of Sciences 1469: 86–104.

Chen, C. et al. 2019. Increasing interannual variability of global vegetation greenness. - Environmental Research Letters 14: 124005.

Chesson, P. 2000. Mechanisms of Maintenance of Species Diversity. - Annual Review of Ecology, Evolution, and Systematics 31: 343–366.

Chesson, P. 2017. AEDT: A new concept for ecological dynamics in the ever-changing world. - PLoS Biology 15: 1–13.

Chesson, P. and Warner, R. 1981. Environmental variability promotes coexistence in lottery competitive systems. - American Naturalist 117: 923–943.

Chesson, P. and Huntly, N. 1997. The Roles of Harsh and Fluctuating Conditions in the Dynamics of Ecological Communities. - American Naturalist 150: 519–553.

Chesson, P. and Kuang, J. J. 2008. The interaction between predation and competition. - Nature 456: 235–238.

Clark, G. F. and Johnston, E. L. 2011. Temporal change in the diversity-invasibility relationship in the presence of a disturbance regime. - Ecology Letters 14: 52–57.

Clark, J. S. et al. 2010. High-dimensional coexistence based on individual variation: A synthesis of evidence. - Ecological Monographs 80: 569–608.

Cohen, J. M. et al. 2020. Avian responses to extreme weather across functional traits and temporal scales. - Global Change Biology 26: 4240–4250.

Coleman, M. A. and Wernberg, T. 2020. The Silver Lining of Extreme Events. - Trends in Ecology & Evolution 35: 1065–1067.

Compagnoni, A. et al. 2021. Herbaceous perennial plants with short generation time have stronger responses to climate anomalies than those with longer generation time. - Nature Communications 12: 1824.

Côté, I. M. et al. 2016. Interactions among ecosystem stressors and their importance in conservation. - Proceedings of the Royal Society B: Biological Sciences 283: 20152592.

Coulson, T. et al. 2004. Skeletons, noise and population growth: the end of an old debate? - Trends in Ecology & Evolution 19: 359–364.

Courchamp, F. et al. 1999. Inverse density dependence and the Allee effect. - Trends in Ecology & Evolution 14: 405–410.

Danino, M. et al. 2018. Stability of two-species communities: Drift, environmental stochasticity, storage effect and selection. - Theoretical Population Biology 119: 57–71.

Davis, A. J. et al. 1998. Individualistic species responses invalidate simple physiological models of community dynamics under global environmental change. - Journal of Animal Ecology 67: 600–612.

Davis, M. A. et al. 2000. Fluctuating resources in plant communities: A general theory of invasibility. - Journal of Ecology 88: 528–534.

De Palma, A. et al. 2017. Large reorganizations in butterfly communities during an extreme weather event. - Ecography 40: 577–585.

Dean, A. M. and Shnerb, N. M. 2020. Stochasticity-induced stabilization in ecology and evolution: a new synthesis. - Ecology 101: 1–14.

Dee, L. E. et al. 2020. Temperature variability alters the stability and thresholds for collapse of interacting species. - Philosophical Transactions of the Royal Society B: Biological Sciences 375: 20190457.

Dennis, B. 2002. Allee Effects in Stochastic Populations. - Oikos 96: 389–401.

Diez, J. M. et al. 2012. Will extreme climatic events facilitate biological invasions? - Frontiers in Ecology and the Environment 10: 249–257.

Drake, J. M. 2005. Population effects of increased climate variation. - Proceedings of the Royal Society B: Biological Sciences 272: 1823–1827.

Drake, J. M. and Lodge, D. M. 2006. Allee effects, propagule pressure and the probability of establishment: Risk analysis for biological invasions. - Biological Invasions 8: 365–375.

Early, R. and Keith, S. A. 2019. Geographically variable biotic interactions and implications for species ranges. - Global Ecology and Biogeography 28: 42–53.

Ellner, S. P. et al. 2019. An expanded modern coexistence theory for empirical applications (J Metcalf, Ed.). - Ecology Letters 22: 3–18.

Elton, C. S. 1958. The ecology of invasions by animals and plants. - Methuen, London.

Ettinger, A. and HilleRisLambers, J. 2017. Competition and facilitation may lead to asymmetric range shift dynamics with climate change. - Global Change Biology 23: 3921–3933.

Felton, A. J. and Smith, M. D. 2017. Integrating plant ecological responses to climate extremes from individual to ecosystem levels. - Philosophical Transactions of the Royal Society B: Biological Sciences 372: 20160142.

Fey, S. B. and Vasseur, D. A. 2016. Thermal variability alters the impact of climate warming on consumer-resource systems. - Ecology 97: 1690–1699.

Fox, J. W. 2013. The intermediate disturbance hypothesis should be abandoned. - Trends in Ecology and Evolution 28: 86–92.

Fung, T. et al. 2020. Temporal population variability in local forest communities has mixed effects on tree species richness across a latitudinal gradient. - Ecology Letters 23: 160–171.

Germain, R. M. et al. 2018. The ‘filtering’ metaphor revisited: Competition and environment jointly structure invasibility and coexistence. - Biology Letters 14: 20180460.

Gilman, S. E. et al. 2010. A framework for community interactions under climate change. - Trends in Ecology & Evolution 25: 325–331.

Godsoe, W. et al. 2017. Integrating Biogeography with Contemporary Niche Theory. - Trends in ecology & evolution 32: 488–499.

Godsoe, W. et al. 2018. Which Coexistence Mechanisms Should Biogeographers Quantify? A Reply to Alexander et al. - Trends in Ecology and Evolution 33: 145–147.

Grainger, T. N. et al. 2019a. Applying modern coexistence theory to priority effects. - Proceedings of the National Academy of Sciences 116: 6205–6210.

Grainger, T. N. et al. 2019b. The Invasion Criterion: A Common Currency for Ecological Research. - Trends in ecology & evolution 34: 925–935.

Gravel, D. et al. 2011. Species coexistence in a variable world. - Ecology Letters 14: 828–839.

Gudmundson, S. et al. 2015. Environmental variability uncovers disruptive effects of species’ interactions on population dynamics. - Proceedings of the Royal Society B: Biological Sciences 282: 20151126.

Hallett, L. M. et al. 2019. Rainfall variability maintains grass-forb species coexistence. - Ecology Letters 22: 1658–1667.

Harley, C. D. G. and Paine, R. T. 2009. Contingencies and compounded rare perturbations dictate sudden distributional shifts during periods of gradual climate change. - Proceedings of the National Academy of Sciences of the United States of America 106: 11172–11176.

Harris, R. M. B. et al. 2020. Biological responses to extreme weather events are detectable but difficult to formally attribute to anthropogenic climate change. - Sci Rep 10: 14067.

Hart, S. P. et al./person-group>. 2018. How to quantify competitive ability (H de Kroon, Ed.). - Journal of Ecology 106: 1902–1909.

He, Q. et al. 2013. Global shifts towards positive species interactions with increasing environmental stress. - Ecology Letters 16: 695–706.

Heino, M. et al. 2000. Extinction risk under coloured environmental noise. - Ecography 23: 177–184.

Holt, R. D. and Keitt, T. H. 2005. Species’ borders: A unifying theme in ecology. - Oikos 108: 3–6.

Holt, G. and Chesson, P. 2014. Variation in moisture duration as a driver of coexistence by the storage effect in desert annual plants. - Theoretical Population Biology 92: 36–50.

Huntingford, C. et al. 2013. No increase in global temperature variability despite changing regional patterns. - Nature 500: 327–330.

Hutchinson, G. E. 1961. The paradox of the plankton. - American Naturalist 95: 137–145.

Hutchison, C. et al. 2020. Seasonal food webs with migrations: multi-season models reveal indirect species interactions in the Canadian Arctic tundra. - Philosophical Transactions of the Royal Society A: Mathematical, Physical and Engineering Sciences 378: 20190354.

Janzen, D. H. 1967. Why Mountain Passes are Higher in the Tropics. - The American Naturalist 101: 233–249.

Jongejans, E. et al. 2010. Plant populations track rather than buffer climate fluctuations. - Ecology Letters 13: 736–743.

Kahilainen, A. et al. 2018. Metapopulation dynamics in a changing climate: Increasing spatial synchrony in weather conditions drives metapopulation synchrony of a butterfly inhabiting a fragmented landscape. - Global Change Biology 24: 4316–4329.

Ke, P.-J. and Letten, A. D. 2018. Coexistence theory and the frequency-dependence of priority effects. - Nature Ecology & Evolution 2: 1691–1695.

Keitt, T. H. et al. 2001. Allee effects, invasion pinning, and species’ borders. - American Naturalist 157: 203–216.

Kennedy, J. P. et al. 2020. Hurricanes overcome migration lag and shape intraspecific genetic variation beyond a poleward mangrove range limit. - Molecular Ecology 29: 2583–2597.

Kraft, N. J. B. et al. 2015. Community assembly, coexistence and the environmental filtering metaphor. - Functional Ecology 29: 592–599.

Kramer, A. M. et al. 2018. Editorial: Allee effects in ecology and evolution. - Journal of Animal Ecology 87: 7–10.

Lande, R. 1993. Risks of Population Extinction from Demographic and Environmental Stochasticity and Random Catastrophes. - The American Naturalist 142: 911–927.

Lawson, C. R. et al. 2015. Environmental variation and population responses to global change. - Ecology Letters 18: 724–736.

Le Coeur, C. et al. 2021. Population responses to observed climate variability across multiple organismal groups. - Oikos: oik.07371.

Legault, G. et al. 2020. Interspecific competition slows range expansion and shapes range boundaries. - Proceedings of the National Academy of Sciences 117: 26854–26860.

Leibold, M. A. et al. 2021. The internal structure of metacommunities. - Oikos in press.

Lenanton, R. C. J. et al. 2017. Potential influence of a marine heatwave on range extensions of tropical fishes in the eastern Indian Ocean—Invaluable contributions from amateur observers. - Regional Studies in Marine Science 13: 19–31.

Lenoir, J. et al. 2020. Species better track climate warming in the oceans than on land. - Nature Ecology & Evolution 4: 1044–1059.

Letten, A. D. et al. 2018. Species coexistence through simultaneous fluctuation-dependent mechanisms. - Proceedings of the National Academy of Sciences of the United States of America 115: 6745–6750.

Levine, J. M. and Rees, M. 2004. Effects of temporal variability on rare plant persistence in annual systems. - American Naturalist 164: 350–363.

Levins, R. 1979. Coexistence in a Variable Environment. - American Naturalist 114: 765–783.

Louthan, A. M. et al. 2015. Where and When do Species Interactions Set Range Limits? - Trends in Ecology & Evolution 30: 780–792.

Louthan, A. M. et al. 2021. Climate sensitivity across latitude: scaling physiology to communities. - Trends in Ecology & Evolution 36: 931–942.

Lyberger, K. et al. 2021. Is evolution in response to extreme events good for population persistence? - The American Naturalist: 714419.

MacDougall, A. S. et al. 2009. Plant invasions and the niche. - Journal of Ecology 97: 609–615.

Maxwell, S. L. et al. 2019. Conservation implications of ecological responses to extreme weather and climate events. - Diversity and Distributions 25: 613–625.

Medeiros, L. P. et al. 2020. Observed ecological communities are formed by species combinations that are among the most likely to persist under changing environments. - American Naturalist 197: E17–E29.

Melbourne, B. A. and Hastings, A. 2008. Extinction risk depends strongly on factors contributing to stochasticity. - Nature 454: 100–103.

Melbourne, B. A. et al. 2007. Invasion in a heterogeneous world: Resistance, coexistence or hostile takeover? - Ecology Letters 10: 77–94.

Miller, T. E. X. et al. 2020. Eco-evolutionary dynamics of range expansion. - Ecology 101: e03139.

Miller, A. D. et al. 2021. How disturbance history alters invasion success: biotic legacies and regime change (E Seabloom, Ed.). - Ecology Letters 24: 687–697.

Mills, S. C. et al. 2017. European butterfly populations vary in sensitivity to weather across their geographical ranges. - Global Ecology and Biogeography 26: 1374–1385.

Myers-Smith, I. H. et al. 2015. Climate sensitivity of shrub growth across the tundra biome. - Nature Climate Change 5: 887–891.

Nadeau, C. P. and Urban, M. C. 2019. Eco-evolution on the edge during climate change. - Ecography 42: 1280–1297.

Nadeau, C. P. et al. 2017. Climates Past, Present, and Yet-to-Come Shape Climate Change Vulnerabilities. - Trends in Ecology & Evolution 32: 786–800.

O’Brien, E. K. et al. 2017. Testing for local adaptation and evolutionary potential along altitudinal gradients in rainforest Drosophila: beyond laboratory estimates. - Global Change Biology 23: 1847–1860.

Ockendon, N. et al. 2014. Mechanisms underpinning climatic impacts on natural populations: altered species interactions are more important than direct effects. - Global Change Biology 20: 2221–2229.

Oldfather, M. F. et al. 2020. Range edges in heterogeneous landscapes: Integrating geographic scale and climate complexity into range dynamics. - Global Change Biology 26: 1055–1067.

O’Sullivan, J. D. et al. 2019. Metacommunity-scale biodiversity regulation and the self-organised emergence of macroecological patterns. - Ecology Letters 22: 1428–1438.

Palmer, G. et al. 2017. Climate change, climatic variation and extreme biological responses. - Philosophical Transactions of the Royal Society B: Biological Sciences 372: 20160144.

Pande, J. et al. 2019. Mean growth rate when rare is not a reliable metric for persistence of species. - Ecology Letters: ele.13430.

Paquette, A. and Hargreaves, A. L. 2021. Biotic interactions are more often important at species’ warm versus cool range edges. - Ecology Letters 24: 2427–2438.

Parmesan, C. and Yohe, G. 2003. A globally coherent fingerprint of climate change. - Nature 421: 37–42.

Pearson, R. G. and Dawson, T. P. 2003. Predicting the impacts of climate change on the distribution of species: are bioclimate envelope models useful? - Global Ecology and Biogeography 12: 361–371.

Pecl, G. T. et al. 2017. Biodiversity redistribution under climate change: Impacts on ecosystems and human well-being. - Science 355: eaai9214.

Petchey, O. L. et al. 1997. Effects on population persistence: the interaction between environmental noise colour, intraspecific competition and space. - Proceedings of the Royal Society B: Biological Sciences 264: 1841–1847.

Pinto, S. M. and Ortega, Y. K. 2016. Native species richness buffers invader impact in undisturbed but not disturbed grassland assemblages. - Biological Invasions 18: 3193–3204.

Platts, P. J. et al. 2019. Habitat availability explains variation in climate-driven range shifts across multiple taxonomic groups. - Scientific Reports 9: 1–10.

Rehage, J. S. et al. 2016. Knocking back invasions: Variable resistance and resilience to multiple cold spells in native vs. nonnative fishes. - Ecosphere 7: 1–13.

Renton, M. et al. 2014. How will climate variability interact with long-term climate change to affect the persistence of plant species in fragmented landscapes? - Environmental Conservation 41: 110–121.

Román-Palacios, C. and Wiens, J. J. 2020. Recent responses to climate change reveal the drivers of species extinction and survival. - Proceedings of the National Academy of Sciences of the United States of America 117: 4211–4217.

Ruel, J. J. and Ayres, M. P. 1999. Jensen’s inequality predicts effects of environmental variation. - Trends in Ecology and Evolution 14: 361–366.

Ryser, R. et al. 2021. Landscape heterogeneity buffers biodiversity of simulated meta-food-webs under global change through rescue and drainage effects. - Nature Communications 12: 4716.

Schreiber, S. J. 2021. Positively and negatively autocorrelated environmental fluctuations have opposing effects on species coexistence. - American Naturalist 197: 405–414.

Schreiber, S. J. et al. 2019. When rarity has costs: coexistence under positive frequency-dependence and environmental stochasticity. - Ecology 100: e02664.

Schultz, E. L. et al. Climate-driven, but dynamic and complex? A reconciliation of competing hypotheses for species’ distributions. - Ecology Letters in press.

Seddon, A. W. R. et al. 2016. Sensitivity of global terrestrial ecosystems to climate variability. - Nature 531: 229–232.

Sexton, J. P. et al. 2009. Evolution and ecology of species range limits. - Annual Review of Ecology, Evolution, and Systematics 40: 415–436.

Shea, K. and Chesson, P. 2002. Community ecology theory as a framework for biological invasions. - Trends in Ecology & Evolution 17: 170–176.

Shoemaker, L. G. et al. 2020a. Integrating the underlying structure of stochasticity into community ecology. - Ecology 101: e02652.

Shoemaker, L. G. et al./person-group>. 2020b. Quantifying the relative importance of variation in predation and the environment for species coexistence (R Snyder, Ed.). - Ecology Letters 23: 939–950.

Sirén, A. P. K. and Morelli, T. L. 2020. Interactive range-limit theory (iRLT): An extension for predicting range shifts. - Journal of Animal Ecology 89: 940–954.

Smale, D. A. and Wernberg, T. 2013. Extreme climatic event drives range contraction of a habitat-forming species. - Proceedings of the Royal Society B: Biological Sciences 280: 20122829.

Smith, M. D. 2011. An ecological perspective on extreme climatic events: A synthetic definition and framework to guide future research. - Journal of Ecology 99: 656–663.

Snyder, R. E. 2008. When does environmental variation most influence species coexistence? - Theoretical Ecology 1: 129–139.

Song, C. et al. 2019. On the Consequences of the Interdependence of Stabilizing and Equalizing Mechanisms. - The American Naturalist 194: 627–639.

Soroye, P. et al. 2020. Climate change contributes to widespread declines among bumble bees across continents. - Science 367: 685–688.

Svenning, J. C. et al. 2014. The influence of interspecific interactions on species range expansion rates. - Ecography 37: 1198–1209.

Swain, D. L. et al. 2018. Increasing precipitation volatility in twenty-first-century California. - Nature Climate Change 8: 427–433.

Taheri, S. et al. 2021. Discriminating climate, land-cover and random effects on species range dynamics. - Global Change Biology 27: 1309–1317.

Taylor, C. M. and Hastings, A. 2005. Allee effects in biological invasions. - Ecology Letters 8: 895–908.

Terry, J. C. D. et al. 2021. Natural enemies have inconsistent impacts on the coexistence of competing species. - Journal of Animal Ecology 90: 2277–2288.

Thompson, P. L. and Gonzalez, A. 2017. Dispersal governs the reorganization of ecological networks under environmental change. - Nature Ecology & Evolution 1: 0162.

Thompson, P. L. and Fronhofer, E. A. 2019. The conflict between adaptation and dispersal for maintaining biodiversity in changing environments. - Proceedings of the National Academy of Sciences of the United States of America 116: 21061–21067.

Thompson, R. M. et al. 2013. Means and extremes: Building variability into community-level climate change experiments. - Ecology Letters 16: 799–806.

Thompson, P. L. et al.. 2020. A process-based metacommunity framework linking local and regional scale community ecology (V Calcagno, Ed.). - Ecology Letters 23: 1314–1329.

Tingley, M. W. et al. 2012. The push and pull of climate change causes heterogeneous shifts in avian elevational ranges. - Global Change Biology 18: 3279–3290.

Tucker, C. M. and Cadotte, M. W. 2013. Fire variability, as well as frequency, can explain coexistence between seeder and resprouter life histories. - Journal of Applied Ecology 50: 594–602.

Tucker, C. M. and Fukami, T. 2014. Environmental variability counteracts priority effects to facilitate species coexistence: evidence from nectar microbes. - Proceedings of the Royal Society B: Biological Sciences 281: 20132637.

Turner, M. G. et al. 2020. Climate change, ecosystems and abrupt change: science priorities. - Philosophical Transactions of the Royal Society B: Biological Sciences 375: 20190105.

Twiname, S. et al. 2020. A cross-scale framework to support a mechanistic understanding and modelling of marine climate-driven species redistribution, from individuals to communities. - Ecography: –15.

Tylianakis, J. M. and Morris, R. J. 2017. Ecological Networks Across Environmental Gradients. - Annual Review of Ecology, Evolution, and Systematics 48: 25–48.

Urban, M. C. 2020. Climate-tracking species are not invasive. - Nature Climate Change 10: 382–384.

Urban, M. C. and De Meester, L. 2009. Community monopolization: Local adaptation enhances priority effects in an evolving metacommunity. - Proceedings of the Royal Society B: Biological Sciences 276: 4129–4138.

Urban, M. C. et al. 2012. On a collision course: competition and dispersal differences create no-analogue communities and cause extinctions during climate change. - Proceedings of the Royal Society B: Biological Sciences 279: 2072–2080.

Urban, M. C. et al. 2013. Moving forward: dispersal and species interactions determine biotic responses to climate change. - Annals of the New York Academy of Sciences 1297: 44–60.

Urban, M. C. et al. 2016. Improving the forecast for biodiversity under climate change. - Science 353: aad8466.

Uricchio, L. H. et al. 2019. Priority effects and nonhierarchical competition shape species composition in a complex grassland community. - American Naturalist 193: 213–226.

Usinowicz, J. and Levine, J. M. 2018. Species persistence under climate change: a geographical scale coexistence problem. - Ecology Letters 21: 1589–1603.

Usinowicz, J. and Levine, J. M. 2021. Climate-driven range shifts reduce persistence of competitors in a perennial plant community. - Global Change Biology: gcb.15517.

Usinowicz, J. et al. 2017. Temporal coexistence mechanisms contribute to the latitudinal gradient in forest diversity. - Nature 550: 105–108.

Vasseur, D. a and Fox, J. W. 2007. Environmental fluctuations can stabilize food web dynamics by increasing synchrony. - Ecology Letters 10: 1066–74.

Vasseur, D. A. et al. 2014. Increased temperature variation poses a greater risk to species than climate warming. - Proceedings of the Royal Society B: Biological Sciences 281: 20132612.

Vázquez, D. P. et al. 2017. Ecological and evolutionary impacts of changing climatic variability. - Biological Reviews 92: 22–42.

Vilà-Cabrera, A. et al. 2019. Refining predictions of population decline at species’ rear edges. - Global Change Biology 25: 1549–1560.

Wallingford, P. D. et al. 2020. Adjusting the lens of invasion biology to focus on the impacts of climate-driven range shifts. - Nature Climate Change 10: 398–405.

Wernberg, T. et al. 2013. An extreme climatic event alters marine ecosystem structure in a global biodiversity hotspot. - Nature Climate Change 3: 78–82.

Williams, C. K. et al. 2003. Population dynamics across geographical ranges: Time-series analyses of three small game species. - Ecology 84: 2654–2667.

Yackulic, C. B. 2017. Competitive exclusion over broad spatial extents is a slow process: evidence and implications for species distribution modeling. - Ecography 40: 305–313.

Zander, A. et al. 2017. Effects of temperature variability on community structure in a natural microbial food web. - Global Change Biology 23: 56–67.

Zepeda, V. and Martorell, C. 2019. Fluctuation-independent niche differentiation and relative non- linearity drive coexistence in a species-rich grassland. - Ecology 100: 1–10.

