## Supplementary material for "Synthesising the multiple impacts of climatic variability on community responses to climate change": SI Text

### Supplementary information for 'Synthesising the multiple impacts of environmental variability on species responses to climate change'

**4. Session Information** **15**

This is a dynamically generated document using R Markdown. Source code (.rmd) for this document is redacted for blind review, but will be available online, allowing complete replication. Code to generate the diagrams in figures 2 and 3 is visible in the .rmd source file.

### 1. Population dynamics model incorporating multiple forms of variability

#### Core Model

Throughout, we use a simple discrete time model that tracks the number of individuals of two competing species ($R$, a resident and $I$, an invader) through time at a single site. In the main text we also refer to these as a 'focal species' and a 'competitor species'. The model is designed to be as simple as possible to illustrate the processes in question.

$$R_{t+1} = \delta R_t + \frac{\lambda_R \omega_{R,t} R_t}{1 + \alpha_{RI}\omega_{I,t} I_t + \alpha_{RR} R_t}$$

$$I_{t+1} = \delta I_t + \frac{\lambda_I \omega_{I,t} I_t}{1 + \alpha_{II} I_t + \alpha_{IR}\omega_{R,t} R_t}$$

In this model, the number of each population in the next generation depends on the carryover rate of the site ($\delta$) and the reproductive output of each species set by a Beverton-Holt response. The performance of each species, in terms of both their maximum fecundity in the absence of competition ($\lambda$) and the competitive pressure they exert on the other species ($\alpha$) varies due to climatic variability ($\omega$). We assume the effects of climate on the performance of both species and their ability to exert competitive pressure on the other are linked - we just use two $\omega$ terms. We do not include climate sensitivity in the intra-specific competition terms for simplicity, since our principal focus is on the dynamics of the invader species at low densities. Parameter descriptions are given in Table S1. The model is extended in the next section to include climate change, demographic stochasticity, immigration and Allee effects.

| Parameter | Meaning |
|---|---|
| $R_t$ | Number of the resident population at time step t |
| $I_t$ | Number of the invading population at time step t |
| $\delta$ | Population carry-over rate |
| $\lambda_s$ | Mean intrinsic reproductive rate of species $s$ |
| $\alpha_{ij}$ | Per-capita competitive impact of $j$ on $i$ |
| $\sigma_s$ | Environmental sensitivity of species $s$ |
| $\omega_{s,t}$ | Performance deviation of species $S$ from mean due to climate at time $t$ |
| $\sigma_s$ | Sensitivity deviation of species $S$ to climate |
| $\rho$ | Correlation between the envrionmental responses of the two species |

**Table S1.** Descriptions of parameters in the core model

#### Weather and species performance

Vectors of weather deviations from mean conditions $E$ for species at each timestep are drawn from a multivariate Gaussian distribution, where the correlation between the weather variables affecting each species is determined by $\rho$:

$$\boldsymbol{E} \sim \mathcal{N}_2(\mu_\omega, \Sigma) \ , \qquad \Sigma = \begin{bmatrix} \sigma_R^2 & \rho\sigma_R\sigma_I \\ \rho\sigma_R\sigma_I & \sigma_I^2 \end{bmatrix}$$

These deviations are translated into performances via a Gaussian-curve shaped envrionmental performance function $f_{EPC}(E)$, scaled such that when $E = 0$, $\omega = 1$:

$$\omega_t = f_{EPC}(E_t) = \exp\left[\eta^2 - (E_t - \eta)^2)\right]$$

This function was chosen to allow a variety of curvatures via an offset parameter $\eta$. Where $\eta$ is small, the curvature is largely negative and moderate amounts of envrionmental variation will lead to a reduction in mean value. Two examples are illustrated in Figure S1. Where $\eta$ is larger, curvature around the average E (0) becomes positive and moderate amounts of variation will result in an increase in mean value. Note however, in both cases the *geometric* mean value (relevant for the long term growth rate calculation) decreases with variance.

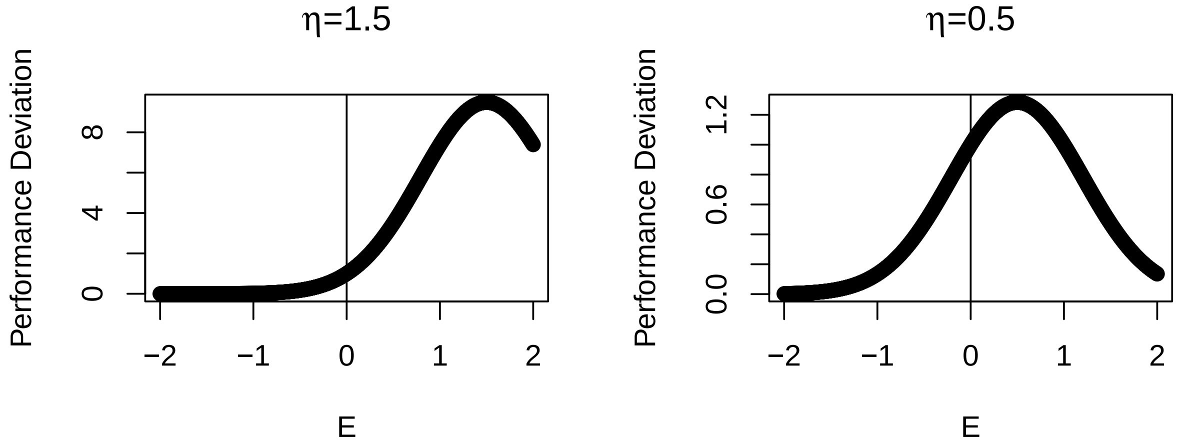

**Figure S1** Demonstration of control of the curvature of the envriomnental performance curve with offset parameter $\eta$.

### 2. Partitioning of invasion growth rate

#### Invasion growth rate

The long-term average growth rate when rare $\overline{r_{inv}}$ describes whether a population would be able to rebound from low levels and hence persist at the site. In a variable environment this rate is determined by multiple interacting mechanisms that can be partitioned to show their direction and relative influence (Ellner et al. 2019). We illustrate here the most direct partitioning, focussing on just the invasion rate of species $I$, and include annotated `R` code in the next section. Here we don't examine coexistence as such, but assume that only the persistence of the focal species, (the 'invader') is in question.

The rate of change in the population using our model (as detailed in the previous section) when the invader is rare ($I_t \to 0$), at a particular time and resident density is given by:

$$\frac{I_{t+1}}{I_t} = \delta + \frac{\lambda_I \omega_{I,t}}{1 + \alpha_{IR}\omega_{R,t}R_t}$$

Since we are using a discrete time model, the growth rate of the invader is calculated on a logarithmic scale:

$$r_{I,t}(\omega_t, R_t) = \ln\left(\frac{I_{t+1}}{I_t}\right)$$

from which the long term mean can be calculated over a large number of time steps:

$$\overline{r_I}(\omega, R) = \frac{1}{t_n}\sum_{t=1}^{t_n} r_{I,t}(\omega_t, R_t)$$

#### Partitioning

The core of the partitioning approach is to adjust this model to successively 'switch-off' different components of variation, and then examine the differences in modeled long-term growth rate. A fixed-baseline model and a fully-variable 'full' model are essential, while the choice of intermediary models will depend on the processes to be considered. Clearly, as the number of fluctuating components rises, there is potentially a very rapid increase in the number of ways this can be done. While in principle building additional models would allow each and every component and interaction term to be determined, there is likely diminishing gains. The selection of which features to change between the models will determine the processes that can be partitioned out - there is no single universal approach, but this flexibility is an advantage. For example, some studies might find it informative to have an even more fundamental baseline, such as the mean of the climate variable under investigation rather than the mean climatic response.

The first step in this process is to determine what the values for all the terms in growth rate model would be without climatic variation or other sources of temporal variation. As discussed above, in our case (by definition), the effect of the mean environment $\overline{\omega} = 1$. The equilibrium resident density can be found directly from $1 - \delta = \lambda_R/(1 - \alpha_{RR}R^*)$ as $R^* = (\lambda_R - 1 + \delta)/(\alpha_{RR}(1 - \delta))$, which we use as our baseline level of residents. Alternatively, the baseline resident population could be set as an average of observed densities, or an average from model simulations $\overline{R}$ to incorporate non-linear responses in the resident population to the environment. Different choices of baseline are valid, but will influence the interpretation of the partition. If there is doubt, multiple choices could be examined and the difference considered as an additional partition.

The next step is to generate a time series of fluctuations in the environment and in the resident population, in the absence of the invader species. This could be derived directly from observations, but more likely to be generated from a statistical model based on longer time series of envrionmental variation. The time series needs to be of sufficient length to capture the full extent of variation patterns.

With these components in hand, the $\overline{r}$ when each aspect of variability is successively 'switched off' can be calculated (Table S2). In our case, we seek to partition the difference between the baseline (fully -fixed) growth rate $\overline{r}_0$ and the overall growth rate $\overline{r}_{full}$ into five parts. This is done by defining based on our understanding of the system (Figure S2) that:

$$\overline{r}_{full} - \overline{r}_0 = \Delta_\lambda + \Delta_\alpha + \Delta_R + \Delta_* + \Delta_{TSE}$$

The contribution of each process term can be directly found from differences between models that incorporate different combinations of terms (Table S3). It follows that the number of alternative models that can be tested determines the number and identity of distinct terms that can be identified. Interaction terms, here $\Delta_*$, capture emergent effects that arise from underlying nonlinearities that cause the impact of multiple processes to differ from a direct sum of their individual effects. As such, they are closer to a correction factor, and do not necessarily have a direct ecological interpretation. In this example we just fit one 'overall' interaction term $\Delta_*$, but in principle (as we have three underlying fluctuating quantities: $\Delta_\lambda, \Delta_\alpha, and \Delta_R$) we could have built additional models to partition this further into emergent effects arising from each pair of processes too. However, in most cases this is unlikely to bring additional ecological insight and there are likely to be limited additional gains for ever finer partitions given the requirements for parameterisation.

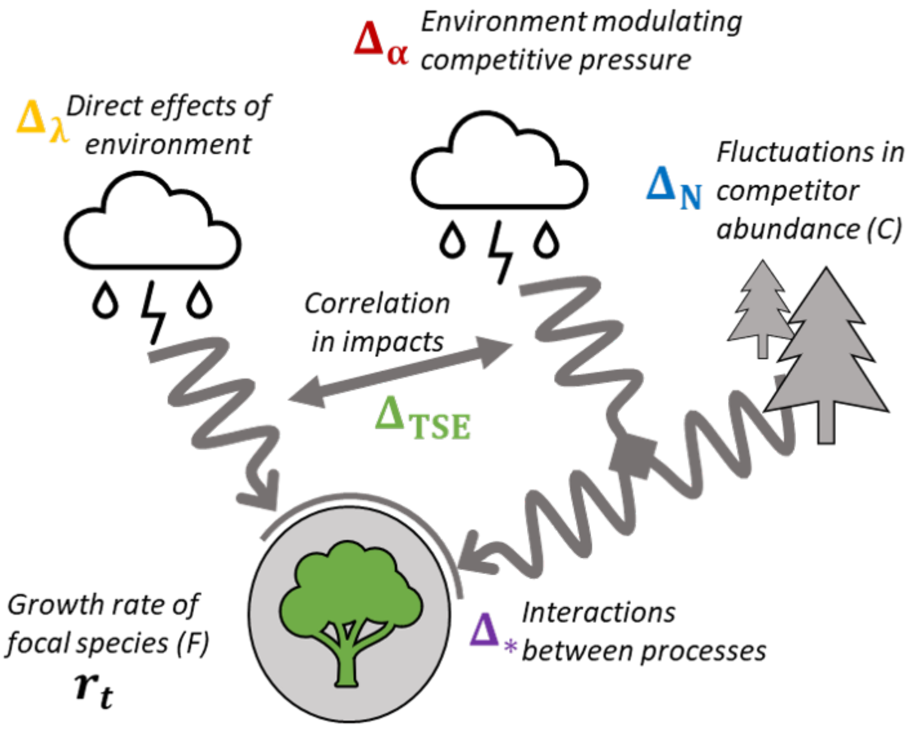

**Figure S2** Summary of processes occurring the system that are included in the model subject to the partitioning.

| Model | Calculation | Description | Partition Components |
|---|---|---|---|
| Baseline | $r_I(\overline{\omega_I}, \overline{\omega_R}, R^*)$ | Baseline model at constant average conditions | $r_0$ |
| A | $\overline{r_I}(\omega_I, \overline{\omega_R}, R^*)$ | Varying invader growth rate with environment | $r_0 + \Delta_\lambda$ |

| Model | Calculation | Description | Partition Components |
|---|---|---|---|
| B | $\overline{r_I}(\overline{\omega_I}, \omega_R, R^*)$ | Varying per-capita competitive impact with environment | $r_0 + \Delta_\alpha$ |
| C | $\overline{r_I}(\overline{\omega_I}, \overline{\omega}_R, R)$ | Varying competitor number | $r_0 + \Delta_R$ |
| D | $\overline{r_I}(\omega_I^\#, \omega_R^\#, R^\#)$ | Varying both environment and competitor number, but without correlation | $\overline{r}_0 + \Delta_\lambda + \Delta_R + \Delta_\alpha + \Delta_*$ |
| Full | $\overline{r_I}(\omega_I, \omega_R, R)$ | Full model, varying both environment and competitor number | $\overline{r}_0 + \Delta_\lambda + \Delta_R + \Delta_\alpha + \Delta_* + \Delta_{TSE}$ |

**Table S2** Description of models built to partition the influence of different sources of variation.

| Component of $\overline{r_{full}}$ | Calculation | Description |
|---|---|---|
| $\overline{r_0}$ | Baseline | Baseline growth rate at constant average conditions |
| $\Delta_\lambda$ | B-$r_0$ | Impact of fluctuating invader productivity |
| $\Delta_\alpha$ | C-$r_0$ | Impact of fluctuating impact of competitors |
| $\Delta_R$ | C-$r_0$ | Impact of fluctuating competitor numbers |
| $\Delta_*$ | D - $\Delta_\lambda - \Delta_R - \Delta_\alpha - r_0$ | Impact of the interaction between fluctuating competitor numbers and environment |
| $\Delta_{TSE}$ | Full - D | Temporal storage effect - Impact of the correlation between species performances in a fluctuating environment |

**Table S3** Description of partitions and how they are calculated from the set of models.

The distinction between Model D and the complete model is worth highlighting, as it is important to identify temporal storage effects. In model D we use a climate variability vector $\omega^\#$ where $\rho = 0$ in order to remove any effects from correlations between the envrionmental responses. This plays a similar role to the randomisation of the order of R to generate $R^\#$ in the examples given in Ellner et al. (2019). In our case, because our environment is not autocorrelated in time, there is no relationship between the *number* of competitors and the weather in each year - here the relevant covariance is between performances. This illustrates how the temporal storage effect derives from covariance between intrinsic growth rates and the *impact of* competitors, not necessarily their number. Where there is a relationship, this could either be considered as a separate partition, or incorporate within one overall 'storage effect' term.

#### Figure 4 examples

To make Figure 4c in the main text, the partitioning described above was carried out on models parameterised as per Table S4. In both cases the envrionmental sensitivity of both species was varied by defining a sequence of average envrionmental sensitivity $\epsilon$. The resident species responded marginally more to the environment than the invader: $\sigma_I = 1.2\epsilon$ and $\sigma_R = 0.8\epsilon$.

| Parameter | Model A | Model B |
|---|---|---|
| $\delta$ | 0.5 | 0.5 |
| $\lambda_R$ | 20 | 16 |
| $\lambda_I$ | 24 | 24 |
| $\rho$ | -0.7 | 0.7 |
| $\alpha_{RR}$ | 0.02 | 0.02 |
| $\alpha_{RI}$ | 0.03 | 0.04 |
| $\alpha_{IR}$ | 0.04 | 0.03 |
| $\eta$ | 1.5 | 0.5 |

**Table S4** Parameters used in the models partitioned in Figure 4c of the main text.

#### Partitioning code

```
## Example model parameters:

Example_Params = list( delta = 0.5,  # Invader dispersal Rate
                       Lam_R = 20,   # Growth rate of Resident
                       Lam_I = 24,   # Growth Rate of Invader
                       W_R_sd = 0.1, # Environment SD for Resident
                       W_I_sd = 0.1, # Environment SD for Invader
                       W_rho = -0.7, # Correlation in environmental effects
                       a_RR = 0.02,  # Resident intraspecific competition
                       a_RI = 0.03 , # Interspecific competition (I on R)
                       a_IR = 0.04,  # Interspecific competition (R on I)
                       eta = 1.5)    # EPC offset from Peak

  ##Function to calculate partitions using the model given a set of parameters.

E_Partition<- function(Params){

  with(Params, {
    n_reps = 10000

    ## Define covariance matrices
    W_SIGMA_corr = matrix(c(W_R_sd^2,
                            W_rho*W_R_sd*W_I_sd,
                            W_rho*W_R_sd*W_I_sd,
                            W_I_sd^2),
                          2,2)

    W_SIGMA_uncorr = matrix(c(W_R_sd^2,
                              0, 0,
                              W_I_sd^2),
                            2,2)

    ## Variable E draws:
    E_mat_full   <-  MASS::mvrnorm(n_reps,   mu = c(0,0),  Sigma = W_SIGMA_corr   )
    E_mat_uncorr<-  MASS::mvrnorm(n_reps,   mu = c(0,0),  Sigma = W_SIGMA_uncorr   )
    E_mat_mean   <-  matrix(c(0,0), nrow = 1)

    ## Convert into \omegas using gaussian curve EPC function
    EPC<- function(WeatherDraws, OffsetFromPeak){
      return(exp( OffsetFromPeak^2-(WeatherDraws -OffsetFromPeak)^2 ))
    }

    W_mat_full   <-   EPC(E_mat_full, eta)
    W_mat_uncorr<-   EPC(E_mat_uncorr, eta)
    W_mat_mean   <-   EPC(E_mat_mean , eta)

    ## Vary only one direct effect of environment
    W_mat_FixR_VarI=   W_mat_FixI_VarR = W_mat_uncorr
    W_mat_FixI_VarR[,2]<- W_mat_mean[2]
    W_mat_FixR_VarI[,1]<- W_mat_mean[1]
```

```
##### Calculating resident fluctuations without invader
R_fluc = rep( NA, n_reps)  # rows = time, cols = space (here 1)
R_fluc[1] <- 10 # initialize

for(t in 1:(n_reps-1)){
  R_fluc[t+1]  =  R_fluc[t]*(delta + Lam_R*W_mat_full[t,1]/(1+a_RR*R_fluc[t]))
}

### Take off a 'burn-in' period
ToUse = (n_reps/10):n_reps
R_var <- R_fluc[ToUse]
W_mat_uncorr<-  W_mat_uncorr[ToUse,]
W_mat_full<-  W_mat_full[ToUse, ]

## resident 'mean' value at equilibrium assuming omega = 1
## R* =
R_mean = (Lam_R-1+delta)/ (a_RR*(1-delta))

## Finding invasion growth rates in different situations;
FindLogGrowthRate<- function( W_Mat, Competitors, Params ){
  with(Params, {
    r_vec<-  delta+ (Lam_I*W_Mat[,2]/( 1+a_IR*Competitors*W_Mat[,1])  )
    return(log(r_vec))
  })
}

## No variation baseline
Zero_r_I =   FindLogGrowthRate(W_mat_mean, R_mean, Params)

## Variation in direct envrionmental effects on invader
A_r_I = FindLogGrowthRate(W_mat_FixR_VarI, R_mean, Params)

## Variation in envrionmental control of competition
B_r_I = FindLogGrowthRate(W_mat_FixI_VarR, R_mean, Params)

### B Variation in number of competitors
C_r_I =   FindLogGrowthRate( W_mat_mean, R_var, Params)

## D Variation in growth rates  and competitors, but not correlated
D_r_I  =  FindLogGrowthRate(W_mat_uncorr,R_var, Params)
## Full model
Full_r_I  =  FindLogGrowthRate(W_mat_full,R_var, Params)

## Take averages of log(r)
A_r_I_mean = mean(A_r_I)
B_r_I_mean = mean(B_r_I)
C_r_I_mean = mean(C_r_I)
D_r_I_mean = mean(D_r_I)
Full_r_I_mean = mean(Full_r_I)

## Calculate partitions
D_Baseline = Zero_r_I
D1_FocalEnvVar = A_r_I_mean - Zero_r_I
```

```
    D2_CompVar = B_r_I_mean - Zero_r_I
    D3_ResiVar = C_r_I_mean - Zero_r_I
    D4_InterVar = D_r_I_mean - (D_Baseline+ D1_FocalEnvVar +D2_CompVar+D3_ResiVar)
    D5_TempStor = Full_r_I_mean - D_r_I_mean

    return(data.frame(Zero_r_I,  A_r_I_mean,  B_r_I_mean,
                      C_r_I_mean,   D_r_I_mean,  Full_r_I_mean,
                      D_Baseline, D1_FocalEnvVar, D2_CompVar,
                      D3_ResiVar,  D4_InterVar, D5_TempStor))})
}

## An Example:
E_Partition(Params = Example_Params)
```

```
##     Zero_r_I A_r_I_mean B_r_I_mean C_r_I_mean D_r_I_mean Full_r_I_mean
## 1 -0.2184079 -0.2107896 -0.2052837 -0.2230557 -0.2044793    -0.1892968
##   D_Baseline D1_FocalEnvVar D2_CompVar   D3_ResiVar  D4_InterVar D5_TempStor
## 1 -0.2184079    0.007618385 0.01312422 -0.004647765 -0.002166153  0.01518247
```

### 3. Interactions between sources of variability

In Figure 6 of the main text, we present some examples of how different sources of variability can interact in their effect on species response to climate change. For this analysis we extend the basic model detailed above to include climate change, immigration, demographic stochasticity and Allee effects.

The model setting is a single site with a resident population. There is incoming dispersal of a potential invader species $I$, but at the start of the trial the invader cannot permanently establish, although it might transiently occur. This is designed to represent a species at its range edge. After a brief establishment phase to equilibriate the resident species, climate change is introduced that makes the environment becomes increasingly more suitable for the invader and less suitable for the resident.

The dynamics are tracked through time. Typically, the invader species is intermittently at densities above 1, before permanently establishing. At the end of the trial the number of time steps between the onset of climate change between which the invader was at a density below 1 is labeled the colonisation time.

In the models we present here we seek to look beyond direct impacts of non-linear averaging in the focal species response to the environment. To do this, as the variance of $\omega$ is varied, we maintain the mean envrionmental response $\overline{\boldsymbol{\omega}}$ fixed at 1 by setting: $\mu_{\omega,i} = -0.5\sigma_i^2$.

#### Model Extensions

##### Climate change

During period of climate change, starting at $t = t_{cc}$, an underlying environment term $\gamma$ increases smoothly from 0 to 1 at the end of the trial ($t_{max}$). Each species response to the underlying environment is controlled by $\beta_s$ via $\omega$:

$$\omega^*_{s,t} = \omega_{s,t} + \beta_s \gamma_t$$

##### Immigration

To represent immigration of the invader into the patch, we introduce an additional time-varying term $d$ to the invader's growth equation:

$$I_{t+1} = \delta I_t + \frac{\lambda_I \omega_{I,t} I_t}{1 + \alpha_{II} I_t + \alpha_{IR} \omega_{R,t} R_t} + d_t$$

A vector of dispersal rates $\boldsymbol{d}$ for each time step is drawn from a log-normal distribution, fixing the resultant mean.

$$log(\boldsymbol{d}) \sim \mathcal{N}(d_\mu - d_\sigma^2/2,\ d_\sigma)$$

##### Allee Effects

To introduce Allee effects for the invader, we instead use a function with a quadratic intraspecific density-dependence component.

$$I_{t+1} = \delta I_t + \frac{\lambda_I \omega_{I,t} I_t^2}{1 + (z + yI_t) I_t + \alpha_{IR} \omega_{R,t} R_t} + d_t$$

Directly comparing functional responses with and without Allee effects is complicated by the impossibility of standardising all other aspects of the multi-species functional response simultaneously. Notwithstanding, we sought chose parameters that given an approximate correspondance between the original models (Fig S2).

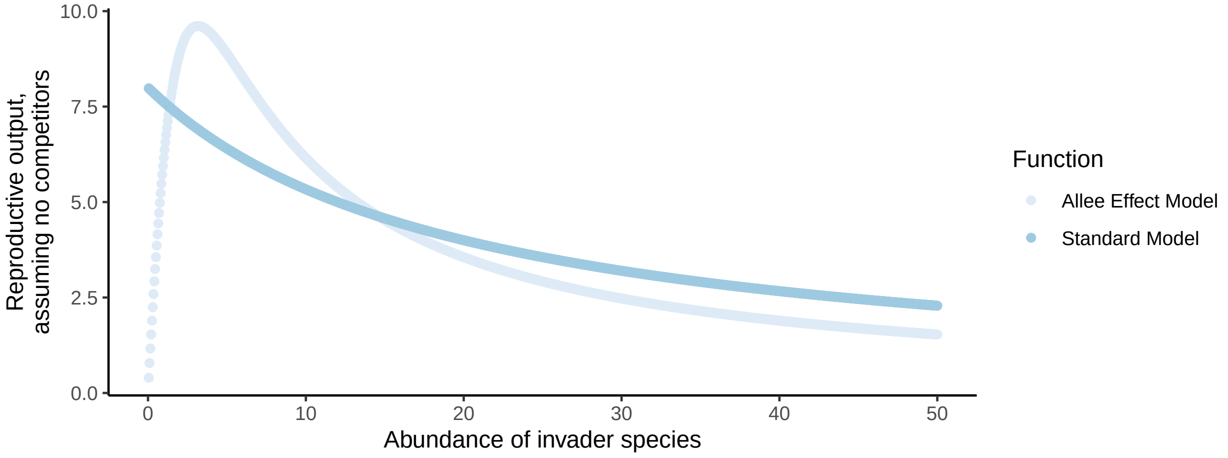

**Figure S3** Comparison between the standard and 'Allee effect' functional responses of the invader using the chosen parameters

**Demographic stochasticity**

To represent demographic stochasticity, the actual number in the next generation is drawn from a Poisson distribution with the expectation value as calculated by the previous equation (Shoemaker et al 2020). This also has the effect of discretising the population densities.

$$I'_{t+1} \sim Pois(I_{t+1}),$$

$$R'_{t+1} \sim Pois(R_{t+1})$$

| Parameter | Meaning |
|---|---|
| $\omega_t^*$ | Effect of climate at time $t$, modified by climate change |
| $z$, $y$ | Invader intra-specific competition coefficient parameters in Allee effect model |
| $d_t$ | Invader incoming dispersal rate |
| $d_\mu$ | (Log) Mean incoming dispersal rate |
| $d_\sigma$ | (Log) Standard deviation of incoming dispersal rate |

**Table S4** Model extension parameter descriptions

#### Figure 6 examples

We use the model to test four interactions between components of variability in terms of impact on colonisation time. Parameter values and key settings are given in Table S5. For clarity in the main text we show average responses as determined by fitting a quadratic curve through 500 trials of each model setup. We show the full results in Figure S4.

| Parameter | a | b | c | d |
|---|---|---|---|---|
| $\delta$ | 0.5 | 0.5 | 0.5 | 0.5 |
| $\lambda_R$ | 20 | 20 | 10 | 9 |
| $\lambda_I$ | 16 | 16 | 6 | 3 |
| $\sigma_R$ | 0.4 | 0.4 | X | 0/X |
| $\sigma_I$ | 0.7 | 0.7 | X | X |
| $\eta$ | 0 | 0 | 0 | 0 |
| $\rho$ | X | 0 | 0 | 0.9 |
| $\alpha_{RR}$ | 0.02 | 0.02 | 0.03 | 0.05 |
| $\alpha_{II}$ | 0.03 | 0.05 | 0.06 | 0.08 |
| $y$, $z$ | - | 0.1,0.2 | - | - |
| $\alpha_{RI}$ | 0.03 | 0.03 | 0.04 | 0.03 |
| $\alpha_{IR}$ | 0.04 | 0.04 | 0.03 | 0.04 |
| $\beta_R$ | -0.1 | -0.1 | -0.1 | -0.1 |
| $\beta_I$ | 0.8 | 0.8 | 0.8 | 0.8 |
| $d_\mu$ | 0.1 | 0.1 | 0.15 | 0.2 |
| $d_\sigma$ | 0.2 | X | 0.01 | 0.01 |
| Allee | FALSE | X | FALSE | FALSE |
| Discretise | FALSE | FALSE | X | FALSE |

**Table S5** Parameter values used in the demonstration of interactions between sources of variability. X indicates that the value was systematically varied.

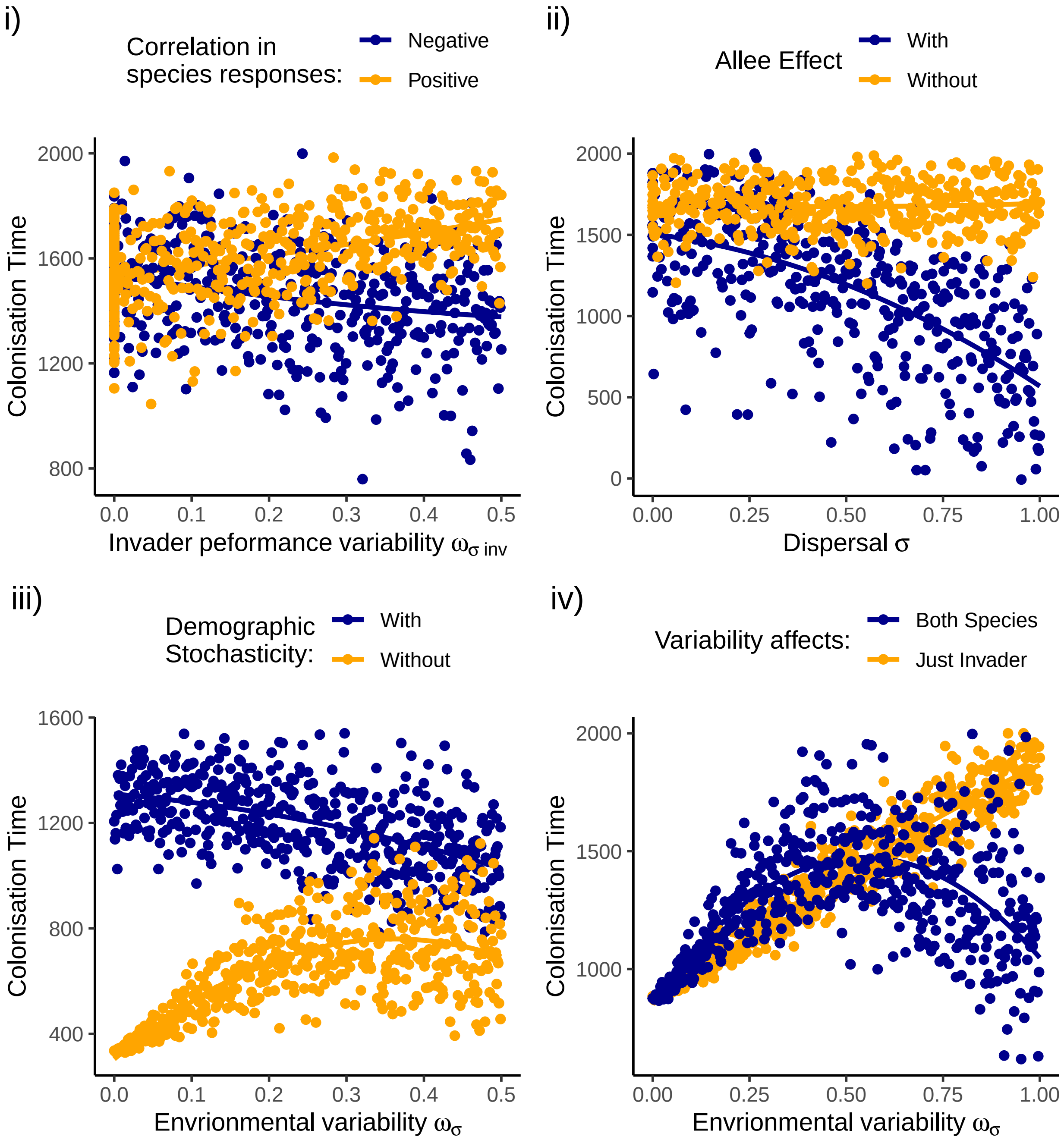

**Figure S4** Raw results underlying the smooth lines presented in Figure 6 of the main text.

### 4. Session Information

```
## R version 4.1.1 (2021-08-10)
## Platform: x86_64-w64-mingw32/x64 (64-bit)
## Running under: Windows 10 x64 (build 19043)
##
## Matrix products: default
##
## locale:
## [1] LC_COLLATE=English_United Kingdom.1252
## [2] LC_CTYPE=English_United Kingdom.1252
## [3] LC_MONETARY=English_United Kingdom.1252
## [4] LC_NUMERIC=C
## [5] LC_TIME=English_United Kingdom.1252
##
## attached base packages:
## [1] stats     graphics  grDevices utils     datasets  methods   base
##
## other attached packages:
##  [1] cowplot_1.1.1   knitr_1.36      forcats_0.5.1   stringr_1.4.0
##  [5] dplyr_1.0.7     purrr_0.3.4     readr_2.0.2     tidyr_1.1.4
##  [9] tibble_3.1.5    ggplot2_3.3.5   tidyverse_1.3.1 MASS_7.3-54
##
## loaded via a namespace (and not attached):
##  [1] Rcpp_1.0.7         lubridate_1.8.0    lattice_0.20-44    assertthat_0.2.1
##  [5] digest_0.6.28      utf8_1.2.2         R6_2.5.1           cellranger_1.1.0
##  [9] backports_1.2.1    reprex_2.0.1       evaluate_0.14      httr_1.4.2
## [13] highr_0.9          pillar_1.6.4       rlang_0.4.11       readxl_1.3.1
## [17] rstudioapi_0.13    Matrix_1.3-4       rmarkdown_2.11     labeling_0.4.2
## [21] splines_4.1.1      munsell_0.5.0      broom_0.7.9        compiler_4.1.1
## [25] modelr_0.1.8       xfun_0.26          pkgconfig_2.0.3    mgcv_1.8-36
## [29] htmltools_0.5.2    tidyselect_1.1.1   viridisLite_0.4.0  fansi_0.5.0
## [33] crayon_1.4.1       tzdb_0.1.2         dbplyr_2.1.1       withr_2.4.2
## [37] grid_4.1.1         nlme_3.1-152       jsonlite_1.7.2     gtable_0.3.0
## [41] lifecycle_1.0.1    DBI_1.1.1          magrittr_2.0.1     scales_1.1.1
## [45] cli_3.0.1          stringi_1.7.5      farver_2.1.0       fs_1.5.0
## [49] xml2_1.3.2         ellipsis_0.3.2     generics_0.1.0     vctrs_0.3.8
## [53] RColorBrewer_1.1-2 tools_4.1.1        glue_1.4.2         hms_1.1.1
## [57] fastmap_1.1.0      yaml_2.2.1         colorspace_2.0-2   isoband_0.2.5
## [61] rvest_1.0.2        haven_2.4.3
```
